# Drinking Motives Synchronize Behavioral and Neural Craving Responses to Alcohol-drinking Videos

**DOI:** 10.64898/2026.07.05.736452

**Authors:** Mina Kwon, Suyeon Song, Hyeonmin Lee, Manjae Kwon, Jung-Seok Choi, Young-Chul Jung, Monica D. Rosenberg, Woo-Young Ahn

## Abstract

Alcohol drinking motives vary among individuals and shape experiences and beliefs about alcohol, influencing the processing of alcohol-related cues. In real-life settings, these cues are contextually rich, amplifying the role of such individualized drinking motives on cue processing. However, previous literature has primarily relied on images of alcohol, which lack contexts and differ significantly from real-life. Here, aiming to investigate real-life craving, we examined the role of alcohol drinking motives in craving in response to naturalistic alcohol-drinking videos. We asked fifty-three problematic alcohol users to speak about their reasons for drinking alcohol to capture unique alcohol drinking motives of each individual. Participants also underwent functional MRI while watching fifteen alcohol-drinking videos, and reported their subjective level of craving and self-relatedness for each video. Behavioral data analysis revealed that individuals with greater alcohol use severity tended to report greater cue-induced craving, but only when they reported that a video was related to themselves. Inter-subject representational similarity analysis showed that participants with similar alcohol drinking motives, reflected in shared drinking reasons and similar self-relatedness to the videos, exhibited synchronized craving-related neural responses during video-watching. Notably, these shared neural processes mediated the link between similar drinking motives and similar self-reported craving levels across participants. Together, our findings highlight the crucial role of alcohol drinking motives in shaping cue-induced alcohol craving, and provide deeper insights into craving in real-world contexts.

## Introduction

People drink alcohol for a variety of reasons: due to habit or stress, for social interactions or emotional comfort, or simply to enjoy the taste of alcohol or food. These individual differences not only shape how we drink, but also how we experience craving: when we feel craving, how strong it is, and whether it leads to actual drinking behavior (1,2).

Craving triggered by drug-related cues is known as cue-induced craving, and its neural mechanisms have been widely investigated using cue exposure paradigms (3). In such paradigms, individuals are typically exposed to images of drugs/alcohol. A robust body of literature has found that individuals with substance use disorders (SUDs) are more likely to feel greater cue-induced craving and exhibit hyperactivation of the brain reward network (4–8). Thereafter, incentive salience theory emerged as one of the well-validated theories that explain addiction: as individuals get addicted to drugs, their reward network hyper-reactivates to drug-related cues, resulting in cue-induced craving (5,9),

A recent functional MRI (fMRI) study examined this theory using a more naturalistic paradigm by showing individuals a drug-themed movie (10). To analyze such naturalistic fMRI data, the authors applied inter-subject correlation (ISC) analysis (11), a data-driven method that identifies brain regions with synchronized (shared) activations across timepoints, to data collected while participants watched the movie. Brain regions exhibiting significant correlations across individuals are interpreted as being strongly modulated by the stimulus (i.e., movie), despite intrinsic and idiosyncratic signals of each individual (12–14). The results suggested that individuals with Heroin Use Disorder showed synchronized activation in the orbitofrontal cortex (OFC), ventromedial and ventrolateral prefrontal cortex, and insula while watching drug-related scenes, which was not observed in healthy controls. Additionally, this drug-biased shared response in the OFC decreased after 15 weeks of treatment, and the reduction was correlated with a decrease in craving. This was the first study to support the incentive salience theory using a naturalistic paradigm. Building on these findings, we applied a similar naturalistic paradigm to alcohol addiction. We further tested whether individual differences in alcohol drinking motives explain such drug-biased neural synchrony in response to real-world cues, beyond the effects of alcohol use severity and self-reported craving.

In real-world settings, alcohol-related cues are rarely presented in isolation but are embedded within rich and dynamic contexts, which amplifies the role of individual differences in processing these cues (15,16). For example, two individuals with similar levels of alcohol use disorder (AUD) symptoms may respond differently to the same cue depending on their memories, preferences, and life experiences. A party scene in a movie might trigger strong craving in someone who typically drinks in social settings but may not affect someone who usually drinks alone.

In this study, we measured these individual differences as alcohol drinking motives, which refer to the broad spectrum of reasons why people drink alcohol. Since Cox and Klinger emphasized the role of motives in alcohol use (17), drinking motives have been recognized as an important factor in understanding alcohol-related behavior (18). Although prior research has shown that drinking motives influence craving responses (19–21), their role in cue-induced craving—particularly in naturalistic settings—remains underexplored. Building on this theoretical background, we empirically tested whether alcohol drinking motives shape how individuals process alcohol-related cues, thereby impacting cue-induced craving in response to naturalistic alcohol-drinking videos. To capture each individual’s drinking motives, we asked participants to speak freely about what drives their alcohol use. Although widely used categories of drinking motives exist (22), we let participants describe their own motives in their own words to capture more personalized and unconstrained expressions of their drinking motives. While this represents a general alcohol drinking motives, not restricted to any predefined models or cues, another important aspect to consider is how well each video (i.e., cue) aligns with an individual’s drinking motives. By asking participants to rate their self-relatedness for each cue—without constraining their responses with predefined constructs such as “related memories,” “preferences,” or “emotional responses”—we captured the extent to which a given cue aligns with an individual’s alcohol drinking motives or alcohol-related experiences.

Together, this study investigated the role of alcohol drinking motives in shaping cue-induced craving within a naturalistic setting. We hypothesized that drinking motives play a critical role in driving cue-induced neural craving, which in turn influences behavioral craving. To test this hypothesis, we first examined whether participants’ behavioral craving could be explained by how well they relate to different alcohol-related video clips. Next, we examined whether participants with similar drinking motives exhibited synchronized activity in a neural signature of craving (neurobiological craving signature, or NCS; (23)) during video-watching. We then tested whether NCS synchrony mediated the link between alcohol drinking motives similarity and behavioral craving similarity. Through this work, we provide a more comprehensive understanding of cue-induced craving in ecologically valid contexts, highlighting the importance of individual drinking motives in alcohol craving in real-world settings.

## Methods

### Participants

Sixty-four individuals participated in the experiment. Participants provided written informed consent and were compensated for their participation. The study was conducted in accordance with the Institutional Review Board at Severance Hospital (IRB No. 4-2023-0162) and at Seoul National University (IRB No. 2302/002-002). Participants first underwent a diagnostic interview based on the Diagnostic and Statistical Manual of Mental disorders, Fifth Edition, Text Revision (DSM-5-TR) to assess AUD and other comorbid mental disorders. Among the sixty-four participants, two did not complete the full experiment, and one was excluded due to invalid responses (i.e., reporting zero craving for all videos). Consequently, data from sixty-one participants were included in the analysis (34 females, age range: 19 to 71, mean age=34.97; SD=10.08; all native Korean speakers).

### Video-watching fMRI Task

Participants completed an fMRI session in which they watched 15 videos displaying naturalistic alcohol consumption scenes. After watching each video, we asked participants to report how much craving for alcohol they felt. We also asked them to report the degree of self-relatedness to the alcohol-drinking scenes in each video clip, that is, how similar the scenes were to their own drinking experiences or motives (***Figure 1A***). An independent behavioral study validated that these videos evoked significantly greater craving than static images of the corresponding beverages (see ***Supplement 1***). Further task details are provided in the ***Supplementary Methods***.

**Figure 1.**
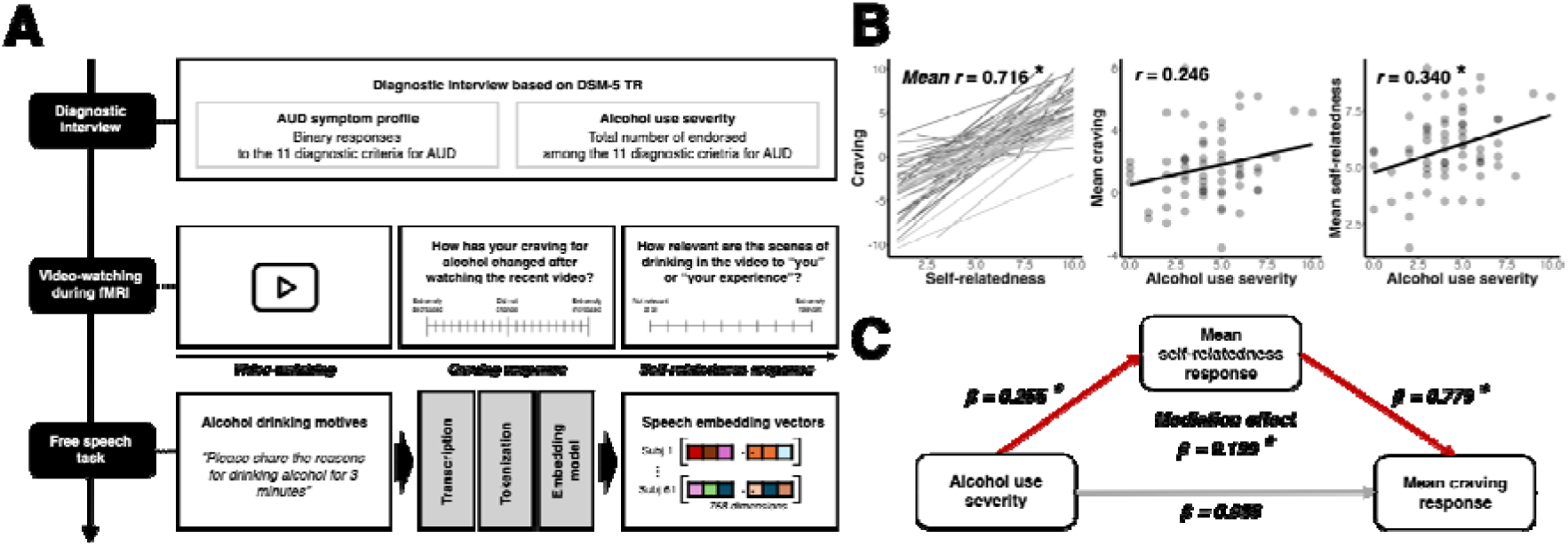
Overview of Study Design and Behavioral Data Analysis. *(A)* Participants first underwent a diagnostic interview for Alcohol Use Disorder (AUD) symptoms based on DSM-5 TR. We used binary responses to the 11 criteria as AUD symptom profiles, and total number of endorsed criteria (i.e., number of AUD symptoms) as alcohol use severity. Within 1-2 weeks, participants completed an fMRI session, during which they watched 15 videos depicting naturalistic alcohol drinking scenes. After watching each video, participants reported their subjective level of alcohol craving (“extremely decreased (−10)” to “extremely increased (10)”). and self-relatedness (“Not relevant at all (0)” to “extremely relevant (10)”), with the order of questions counter-balanced across participants. Following the fMRI session, participants performed a free speech task in which they responded to the prompt: “Please share the reasons for drinking alcohol for 3 minutes.” Their speech was recorded, transcribed, tokenized, and processed using an embedding model to generate 768-dimensional embedding vectors. *(B)* Pairwise relationships among craving, self-relatedness, and alcohol use severity. The left figure shows the correlation between craving and self-relatedness responses for each individual, with each line corresponding to each participant. The correlation coefficient represents the mean value of all individual coefficients. Correlation coefficients were Fisher z-transformed before averaging across participants, and the reported mean *r* reflects the back-transformed group-level average. The middle figure displays the correlation between alcohol use severity and mean craving responses, with each point corresponding to each participant. The right figure displays the correlation between alcohol use severity and mean self-relatedness responses, with each point indicating each participant. *(C)* Mediation model showing that self-relatedness responses fully mediate the relationship between alcohol use severity and craving responses. The coefficient and significance of the mediation effect was estimated based on 10,000 bootstrapped samples. Asterisks indicate statistical significance (*P*<0.05).

### fMRI Image Acquisition and Preprocessing

MRI scanning was performed using a 3.0T Siemens Magnetom TrioTim MRI scanner. Both high-resolution T1-weighted anatomical images and functional echo planar images were obtained. The fMRI data were preprocessed using the standard fMRIPrep pipeline (fMRIPrep version 20.2.0; (24)). After excluding data due to excessive head motion or falling asleep, data from 53 participants were included in the fMRI analysis. The detailed MRI acquisition parameters and the full preprocessing stream are described in the ***Supplementary Methods***.

### Alcohol Drinking Motives

Participants’ alcohol drinking motives were assessed with an open-ended question after the fMRI task. We asked participants to share their alcohol drinking motives for 3 minutes, which was recorded, transcribed to text, and processed into speech embedding vectors using Sentence Transformer (***Figure 1A***). Details on embedding model selection and supplementary analyses using human-rated speech instead of speech embedding vectors are provided in ***Supplement 2***.

### Behavioral Data Analysis

To examine whether an individual’s cue-induced craving could be explained by their self-relatedness ratings and alcohol use severity, we first evaluated the pairwise relationships among these behavioral measures. Next, to compare the explanatory power of self-relatedness and alcohol use severity in predicting craving, we fit a standardized multiple linear mixed-effects model. We also fit a mediation model to test whether self-relatedness could explain the link between alcohol use severity and cue-induced craving. Full details of the analyses are provided in the ***Supplementary Methods***.

### Inter-subject representational similarity analysis (IS-RSA)

We tested whether participants who reported similar craving, relatedness to alcohol videos, and drinking motives showed more synchronous brain activity during video watching, compared to those with more similar AUD symptom profiles or higher alcohol use severity. To do so, we conducted intersubject representational similarity analysis (IS-RSA) to identify brain regions where neural activity was more similar across individuals who exhibited more similar behavior (25). The first step of IS-RSA is to create behavioral similarity matrices (***Figure 2A***). Similarity in craving and self-relatedness was measured with Pearson correlation, drinking motives via cosine similarities between speech embedding vectors, and AUD symptom profiles using Jaccard index. We measured similarity of alcohol use severity scores using the Anna Karenina model (Anna-K). This model tests the hypothesis that individuals with greater or lesser severity exhibit more similar neural responses, whereas individuals on the opposite end of the severity scale are more idiosyncratic (see ***Supplement 3*** for results using the Nearest-neighbor model), in which higher similarity indicates more similar severity). For the neural similarity matrices, based on Shen et al.’s (26) brain parcellation, we created 268 similarity matrices by computing ISC for each of the 268 regions of interest (ROIs) (***Figure 2B***; See ***Figure S14*** for the ISC results). As the final step, we computed the Spearman correlation (i.e., IS-RSA value) between each behavioral similarity matrix and the neural similarity matrices in all ROIs. To determine the significance of the IS-RSA value, we conducted permutation testing with 10,000 permutations. Full details of the significant testing are provided in the ***Supplementary Methods***.

**Figure 2.**
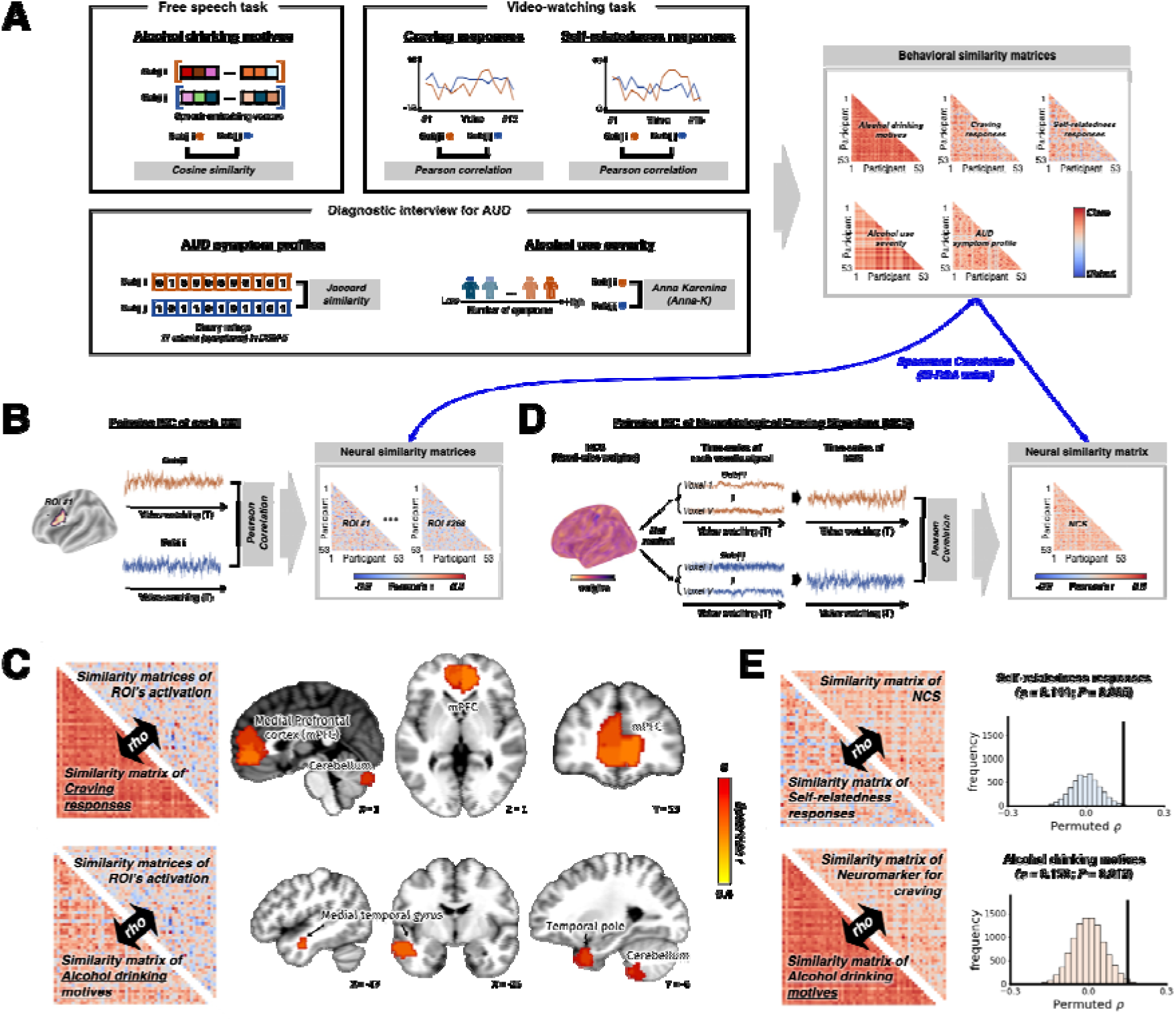
Inter-subject representational similarity analysis (IS-RSA). *(A)* Behavioral data and similarity measures used to create behavioral similarity matrices. Five pairwise (i.e., participant-by-participant) similarity matrices were computed, each representing the similarity between participants based on specific behavioral data: [1] Cosine similarities of speech embedding vectors representing alcohol drinking motives, [2] Pearson correlations between craving responses for each alcohol-drinking video, [3] Pearson correlations between self-relatedness responses for each alcohol-drinking video, [4] Jaccard similarities of binary responses to the 11 diagnostic criteria for Alcohol Use Disorder (AUD), and [5] Similarity in alcohol use severity based on the Anna-Karenina (Anna-K) model. The Anna-K model defines similarity as the average severity rank of a participant pair, inspired by the Tolstoy’s line, “All happy families are alike; each unhappy family is unhappy in its own way.” In this context, the model assumes “all individuals with high [low] severity are alike; each individual with low [high] severity is different in their own way.” *(B)* Creating neural similarity matrices using brain parcellation by Shen et al. (26). Neural similarity matrices were created for each of the 268 regions of interest (ROIs) by computing inter-subject correlation (ISC), which is the Pearson correlation between each ROI’s time-series data for each subject pair. *(C)* Significant results of IS-RSA using 268 neural similarity matrices (B) and the five behavioral similarity matrices (A). Two behavioral similarity matrices—self-relatedness responses and alcohol drinking motives—showed significant Spearman correlations with neural similarity (two-tailed nonparametric test, *P*<0.05, corrected using the Benjamini–Yekutieli (BY) procedure (44) *(D)* Creating a neural similarity matrix using the Neurobiological Craving Signature (NCS; (23)). A single time series of the NCS responses was computed for each participant by taking the dot product of voxel-wise weights of the NCS and voxel-wise activation of each TR. Pearson correlation between these time-series was then used to compute the neural similarity matrix for the NCS. *(E)* Results of IS-RSA using the NCS similarity matrix (D) and the five behavioral similarity matrices (A). Histograms indicate null distribution of 10,000 permuted coefficients, while the black vertical line indicates the IS-RSA value (i.e., Spearman correlation coefficient).

### Evaluating NCS synchrony using IS-RSA

The whole-brain IS-RSA focused on individual brain regions in isolation. Recent work, however, highlighted the importance of widespread patterns of voxel-wise activity in craving and introduced the NCS, a whole-brain map of voxel weights, as a neuromarker of craving (23). Using this pre-defined neural mask, we next directly tested whether “neural craving” is synchronized when individuals share alcohol drinking motives. We calculated a weighted average across brain voxels for each brain image (i.e., for each TR) during video-watching, which resulted in a single NCS score per brain image. When combined across all TRs during video-watching, this yielded a time-series representing fluctuation of the NCS throughout video-watching. Before analyzing the NCS time-series data, we first validated that the NCS was more responsive to alcohol cues than to food cues (***Supplement 4; Figure S8***), as both cues are known to induce craving (27).

Using the time-series of NCS during video-watching, we examined whether participants who reported similar behavior (i.e., craving, relatedness to alcohol videos, drinking motives, AUD symptom profiles, and alcohol use severity) showed more synchronous “neural craving” during video-watching. Since there was a single NCS time series per participant, we created a single neural similarity matrix by computing Pearson correlations between the NCS time series of each pair of participants (***Figure 2D***). We then computed Spearman correlations between this neural similarity matrix and each of the five behavioral similarity matrices (***Figure 2A***). Taking advantage of using the single time series representing neural craving, we further conducted multiple linear regression analyses to evaluate whether these relationships remained significant after controlling for other behavioral similarity measures. The dependent variable was the NCS similarity matrix, and the independent variables were the behavioral similarity matrix with significant correlation from the IS-RSA and the behavioral similarity matrix to control. Full details of the model are provided in the ***Supplementary Methods***. Then, to better understand how shared drinking motives influence neural synchrony and the role of craving responses, we tested a mediation model to determine whether NCS synchrony mediated the relationship between drinking motives and craving. We also tested whether presence of alcohol cues accounted for the mediation effect, given that each video combined alcohol cues, food cues, and neutral contexts. Additional analyses separating the time-series into specific cue conditions are detailed in ***Supplement 5***.

### Assessing Individual Differences in Cue-Driven NCS Synchrony

Finally, as the presence of alcohol could affect NCS synchrony differently for different pairs of individuals, we tested such individual differences. We first fit multiple linear regression models predicting the dynamic ISC of each subject pair using the time-series of alcohol and food cues as predictors (***Figure 3A***). For each model, we obtained beta coefficients for alcohol ({J_alcohol_) and food ({J_food_) cues, and constructed participant-by-participant matrices of {J_alcohol_ and {J_food_, respectively (***Figure 3B***). These matrices represent the degree to which neural synchrony is explained by the presence of cues: a positive beta value indicates that cue presence increases NCS synchrony for that subject pair, while a negative beta value suggests that cue presence decreases NCS synchrony. Next, we tested whether behavioral similarity matrices could explain individual differences in cue-related NCS synchrony (i.e., beta coefficients). We computed Spearman correlations between each beta matrix and the behavioral similarity matrices, respectively (***Figure 3B***). Significance was assessed through permutation testing with 10,000 permutations, consistent with the IS-RSA procedure. For the neural-behavioral pair with significant associations, we examined the absolute values of the beta coefficients. We divided behavioral similarity scores into four quantiles, and for each quantile, we tested whether the mean beta coefficient is significantly different from zero using a one-sample t-test.

**Figure 3.**
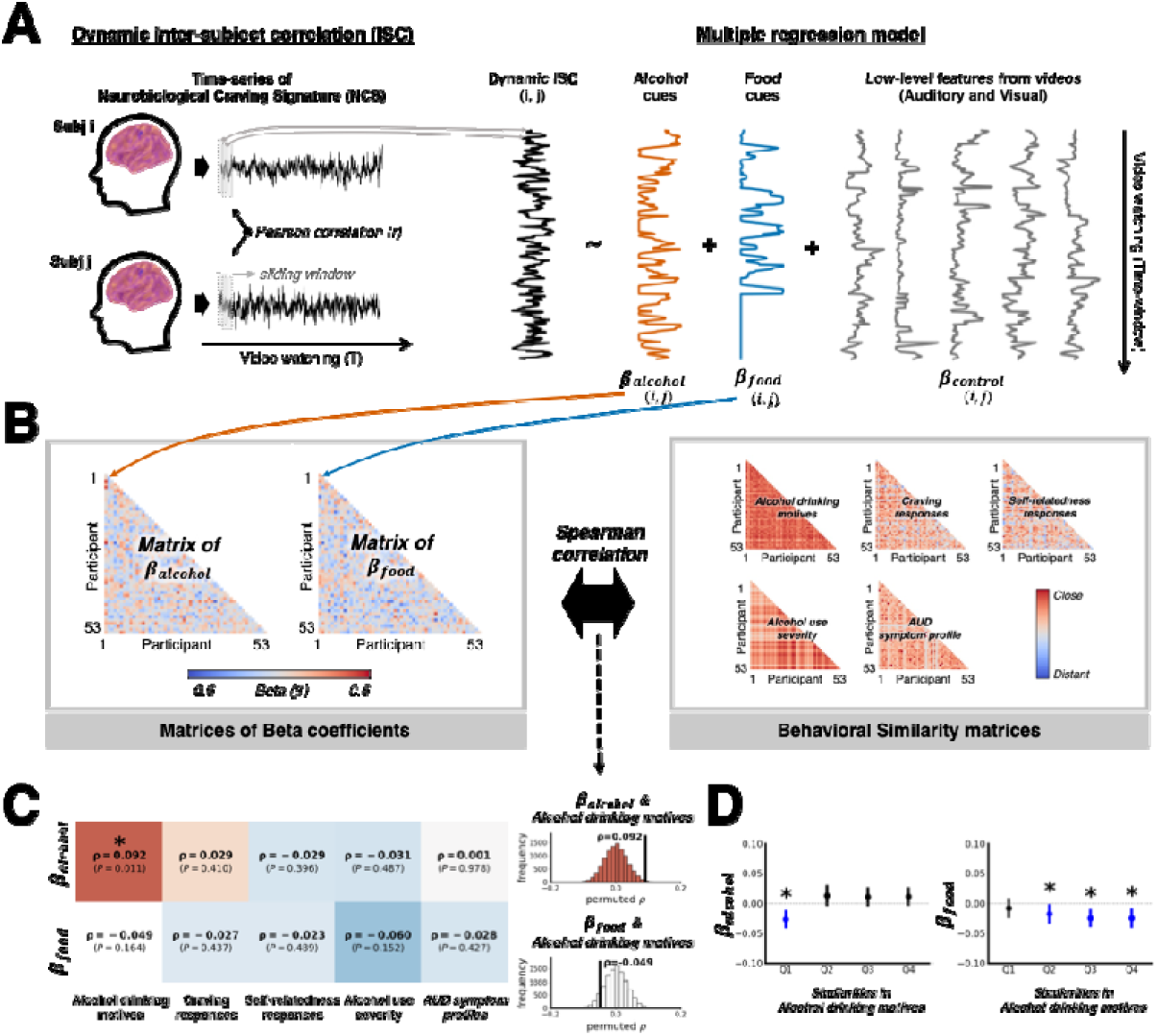
Individual differences in the degree of NCS synchrony explained by alcohol cues. *(A)* Illustration of multiple linear regression model predicting dynamic inter-subject correlation (ISC) of each subject pair, with time series of alcohol and food cue presence as independent variables. A total of 1,378 models were fit. {J_alcohol_ indicates the extent to which alcohol cues explain NCS synchrony in each pair, and {J_food_ indicates the extent to which food cues explain NCS synchrony. *(B)* Spearman correlations between the pairwise beta matrices ({J_alcohol_ and {J_food_) and the pairwise matrices of behavioral similarity (Figure 2A). Higher correlation coefficients between the matrices indicate stronger associations between behavioral similarity and cue-related NCS synchrony. *(C)* Results of the correlation analyses, with statistical significance assessed using permutation testing (10,000 iterations). Histograms show the null distribution of permuted coefficients, with the observed correlation coefficient indicated by black vertical lines. Asterisks denote significant Spearman correlation coefficient (*P*<0.05). *(D)* Analysis of the absolute beta coefficients in relation to similarities in alcohol drinking motives. The X-axis represents similarity values divided into four quantiles (Q1: least similar, Q4: most similar), and the Y-axis displays mean beta coefficients for each quantile. Error bars represent 95% confidence intervals. Asterisks indicate statistically significant mean beta coefficients based on one-sample t-tests.

### Specificity testing

To examine the specificity of the speech embedding vectors of alcohol drinking motives, we repeated the analysis using two additional speech samples collected from the same participants, each matched in duration (3 minutes): speech about quitting drinking (i.e., in response to the prompt “Please share your thoughts about quitting, refraining, or reducing drinking alcohol.”) and control speech (i.e., in response to the prompt “Please share what happened before you came to this laboratory today.”). Furthermore, to evaluate the specificity of the NCS, we repeated all analyses using three published neuromarkers developed to predict constructs other than craving: the Brain Reward Signature (BRS; (28)), the Picture-Induced Negative Emotion Signature (PINES; (29)), and the Neurologic Pain Signature (NPS; (30)). See ***Supplement 6*** for more details.

## Results

### Self-relatedness Mediates the Relationship between Alcohol Use Severity and Cue-induced Craving

We first examined whether an individual’s cue-induced craving could be explained by their self-relatedness ratings and alcohol use severity, by evaluating the one-to-one relationships among these measures. Craving and self-relatedness responses were positively correlated (mean within-subject *r*=0.716, *P*<0.001), suggesting that videos perceived as more self-relevant are more likely to evoke stronger craving. Although participants with more severe alcohol use reported higher craving levels, this relationship was not statistically significant (*r*=0.246, *P=*0.056). However, those with more severe alcohol use tended to report greater self-relatedness (*r*=0.340, *P=*0.007). Then, by fitting the linear mixed-effects model, we found that self-relatedness predicted craving (β=1.121, *P*<0.001) even after controlling for alcohol use severity, while alcohol use severity did not (β=0.148, *P=*0.489). Furthermore, self-relatedness mediated the relationship between alcohol use severity and craving (mediation effect=0.199, 95% CI=[0.050, 0.344], *P*=0.004; ***Figure 1C***). These results suggest that individuals with higher alcohol use severity tend to relate more to alcohol-drinking videos, which in turn explains stronger cue-induced craving.

### Synchronized Brain Regions during Video-watching When Alcohol Drinking Motives are Similar Across Individuals

Given that self-relatedness is more strongly associated with behavioral craving than alcohol use severity, we next investigated the neural underpinnings of this effect. Whole brain IS-RSA results revealed that participants with more similar craving responses (i.e., those who reported more vs. less craving to the same videos) showed greater neural synchrony in three ROIs in the dorsomedial prefrontal cortex (dmPFC; *r=*0.130 ∼ 0.197; corrected *P*=0.004 ∼ 0.006) and posterior cerebellum (*r=*0.113; corrected *P*=0.009) (***Figure 2C***). In addition, participants who reported more similar drinking motives—i.e., those with more similar embeddings of their free speech—showed greater neural synchrony in the middle temporal gyrus, temporal pole, and posterior cerebellum during video-watching. Somewhat surprisingly given behavioral findings, self-relatedness similarity was not significantly associated with neural synchrony. Similarly, AUD symptom profiles and alcohol use severity did not yield any significant ROIs with inter-subject synchrony.

### The Neurobiological Craving Signature (NCS) is synchronized during Video-watching When Alcohol Drinking Motives are Similar

We next applied the NCS to directly test whether “neural craving” is synchronized when individuals share alcohol drinking motives. NCS synchrony was correlated with the similarity of participants’ self-relatedness responses for the videos (p=0.144, *P=*0.006) and drinking motives (p=0.150, *P=*0.012; ***Figure 2E***). No other behavioral measures were significantly associated with NCS synchrony (***Table S1***). Surprisingly, while participants who reported similar craving responses to the drinking videos did not show synchronized NCS responses (p=0.086, *P=*0.156), an interaction between craving similarity and drinking motives similarity significantly predicted NCS synchrony ({J=0.176, *P=*0.026; ***Table S2***). This finding suggests that while similar craving alone did not drive neural synchrony in the NCS, the combination of similar drinking motives and craving responses jointly contributed to NCS synchrony. Overall, individuals with similar alcohol drinking motives—whether specific to the videos (i.e., self-relatedness responses) or in general (i.e., drinking reasons)—tended to exhibit synchronized craving-related neural responses, even when controlling for AUD symptom profiles and alcohol use severity. Furthermore, similarities in craving and drinking motives synergistically predicted synchronized neural craving, whereas craving similarity alone did not account for it. These neural-behavioral patterns were specific to NCS synchrony and alcohol drinking motives, as such patterns were not significant when using alternative speech embeddings or alternative neuromarkers (See ***Supplement 6***).

By fitting the mediation models, we confirmed that NCS synchrony significantly mediated the relationship between synchronized drinking motives and synchronized self-reported craving after video-watching (mediation effect=0.025, 95% CI=[0.009, 0.052], *P*=0.009; ***Figure S5A***). In contrast, the mediation effect was not significant when the independent variable was similarity in self-relatedness responses (β=0.004, 95% CI=[-0.002, 0.011], *P*=0.181; ***Figure S5B***). These results suggest that individuals with similar drinking motives tended to show synchronized NCS responses while watching alcohol-related videos, which in turn corresponded to similar levels of self-reported craving after watching the videos. However, this mediation effect was not observed for the video-specific self-relatedness responses. Furthermore, the mediation effect was significant only when the neural data included timepoints for alcohol cues (See ***Supplement 5 & Figure S6***).

**Diverging Drinking Motives Lead to Idiosyncrasy in the NCS in Response to Alcohol Cues** Lastly, we investigated individual differences in these neural-behavioral patterns by evaluating how cue-driven fluctuations in synchrony varied across participant pairs. While none of the other behavioral measures explained either beta matrix, {J_alcohol_ was significantly correlated with drinking motives similarity (p=0.092, *P=*0.011; ***Figure 3C***), while {J_food_ was not (p=-0.049, *P=*0.164). Further analysis of the absolute beta values revealed that the lowest quantile of similarity in drinking motives had significantly negative beta coefficients (mean of {J_alcohol_ in the first quantile=-0.026, 95% CI=[−0.042, −0.011], *t*=-3.426, *P*<0.001; ***Figure 3D***). This indicates that more dissimilar pairs showed significantly lower {J_alcohol_, thus subject pairs with the most divergent drinking motives exhibited neural desynchrony in the NCS in response to alcohol cues. This relationship was not replicated when using alternative control speech conditions or other neuromarkers (***Supplement 6 & Figure S13***).

## Discussion

Individuals with different alcohol drinking motives may exhibit different patterns of cue-induced craving, particularly within context-rich, naturalistic settings. Our findings demonstrate that alcohol drinking motives play a critical role in cue-induced craving, both in behavioral and neural domains. First, self-relatedness responses to each video, the degree to which each video aligns with one’s alcohol drinking motives, mediated the relationship between alcohol use severity and craving intensity. Furthermore, individuals with shared drinking motives exhibited similar dynamics of the NCS, the predefined neuromarker for craving, while watching naturalistic alcohol-drinking videos, which corresponded to similar levels of behavioral craving. Lastly, individuals with dissimilar drinking motives exhibited more divergent NCS responses when alcohol-related cues were presented. Together, these findings build on prior research that has established a link between greater addiction severity and greater craving (4,6) by highlighting the importance of individual differences in cue-induced craving.

Past studies using static images of alcohol have consistently reported that greater alcohol use severity is associated with heightened level of craving when exposed to alcohol cues. However, when using videos, alcohol use severity alone did not account for craving but was mediated by self-relatedness. We speculate that a key factor underlying this difference is the inclusion of contextual information in the cues. Recent fMRI studies have shown that adding context to alcohol-related images activated brain regions involved in self-referential processing or memory processing (31,32), indicating that adding contextual information engages neurocognitive processes beyond reward processing in cue-induced craving. Consistent with the literature, we also found from IS-RSA that individuals with similar craving responses exhibited synchronized activations in the dmPFC—associated with motivation monitoring and craving regulation (33) and posterior cerebellum, which supports higher-order memory retrieval and cognitive predictions (34,35). Furthermore, individuals sharing similar drinking motives demonstrated synchronized neural dynamics within the default mode network, specifically the middle temporal gyrus and temporal pole (36,37), which is associated with self-referential cognitive states such as autobiographical memory or mind wandering (38). These patterns suggest that shared drinking motives may engage similar neurocognitive mechanisms across individuals during alcohol cue exposure.

IS-RSA results using the NCS further supported this interpretation. Individuals with similar alcohol drinking motives, whether reflected in general drinking reasons or cue-specific self-relatedness, showed more similar neural craving responses during video-watching. These results remained significant even after controlling for similarities in AUD symptom profiles or alcohol use severity. Notably, only general drinking motives were linked to similar craving responses to the cues through the NCS synchrony. We speculate that this divergence may stem from the inherent constraints of video stimuli, which cannot capture the full spectrum of idiosyncratic, real-world experiences, thereby limiting the variance and statistical power of cue-specific self-relatedness ratings. In contrast, general alcohol drinking motives appeared to capture more reliable and individualized information relevant to craving responses. Furthermore, differences in general drinking motives were associated with idiosyncratic neural craving responses to alcohol cues. Crucially, this effect was characterized by desynchronization among dissimilar individuals rather than heightened synchrony among similar ones, potentially reflecting unmeasured individual factors which future research could explore using other types of individual-level data—for example, alcohol-related memories, preferences, or expectations.

We confirmed that this intriguing relationship between the NCS and drinking motives is specific to both constructs. First, drinking motives showed no associations with neural synchrony of alternative neuromarkers (i.e., BRS, PINES, and NPS) that reflects affective and motivational processes closely tied to alcohol addiction. For instance, alcohol use engages reward circuits (39), is often used to cope with negative emotions (19,40), and is associated with heightened pain sensitivity during withdrawal (41–43). While these processes may also be engaged while watching alcohol-drinking videos, only the NCS exhibited neural synchrony based on shared drinking motives, underscoring its specificity. Likewise, despite capturing individualized content, neither the speech about participants’ daily experiences nor the speech about quitting drinking—which contained alcohol-relevant content—explained behavioral and neural craving responses.

Overall, our findings highlight the role of alcohol drinking motives in neural and behavioral craving responses during naturalistic cue exposure. Nonetheless, important questions remain. First, while applying the summary index (i.e., NCS) to the fMRI data provides a direct way to examine “neural craving,” it leaves open the question of the neural mechanisms linking similarity in alcohol drinking motives to similarity in cue-induced craving. Future studies should investigate whether distinct neural mechanisms underlie different motivational subtypes, such as whether memory networks preferentially synchronize among memory-driven drinkers while incentive salience circuits dominate among habitual drinkers. Additionally, although we tested the specificity of the alcohol cues relative to food cues, further testing with videos that do not include alcohol cues may be necessary. Moreover, future studies could assess subjective experiences during video-watching using measures beyond self-relatedness, such as speech or continuous self-report, to compare general versus cue-specific motives more precisely. Finally, future research should replicate our findings using different types of alcohol-related videos, in diverse cultural contexts, with a broader range of participants, and even across different substances to evaluate the generalizability of these results.

## Conclusion

In summary, this study highlights the critical role of individual differences in cue-induced craving, extending the focus beyond addiction severity to alcohol drinking motives. By integrating naturalistic fMRI paradigms with unconstrained free-speech embeddings, we introduce a novel framework for investigating the real-life craving processes. Clinically, these findings underscore the value of personalized treatments that consider *why* and *how* individuals drink rather than relying solely on addiction severity or symptoms.

## Author Contributions

M.K.^1^, S.S, H.M., M.K.^2^, J.-S.C., Y.-C.J. and W.-Y.A. designed and performed research; M.K.^1^, M.D.R., and W.-Y.A. analyzed data and wrote the paper; M.D.R. contributed new reagents/analytic tools.

## AI Disclosure Statement

The authors used ChatGPT-4o and Gemini 3.5 Flash to refine the language and improve readability. After using these tools, the authors reviewed, edited, and approved all final content and take full responsibility for the accuracy of the publication.

## Data Availability

Codes used to run experiments and all codes for the analysis are available at Github, https://github.com/CCS-Lab/project_drinking_motives_craving.

