## supplement for "Drinking Motives Synchronize Behavioral and Neural Craving Responses to Alcohol-drinking Videos"

Kwon *et al.*

**Supplementary Methods**

**Task Description**

During the fMRI session, participants viewed 15 naturalistic video clips that depicted various alcohol consumption scenes, The total task duration was 20.783 minutes, with individual clips ranging from 69 to 106 seconds (mean=89.533, SD=10.162). The selected videos varied in social contexts (7 social drinking, and 8 solo drinking videos), gender of characters (8 male-only, 5 female-only, and 2 mixed-gender videos), type of alcohol (2 soju, 9 beer, 2 wine, 2 makgeolli-drinking videos), and food presence (10 videos with food, and 5 without food). All videos were sourced from Korean observational variety shows, with 10 videos from “*I Live Alone*” and 5 videos from “*My Little Old Boy*.” Following each clip, participants answered two questions evaluating craving and self-relatedness. To control for order effects, the sequence of questions was counterbalanced across participants. Additionally, to minimize order effects of the videos, approximately half of the participants (N=27) watched the videos in reverse order (i.e., starting with the last video and ending with the first).

**Preprocessing fMRI data**

Functional images were acquired using a T2*-weighted gradient echo-planar imaging (EPI) sequence (repetition time [TR]=2,000 ms, echo time [TE]=20 ms, voxel size=3.0 mm, slice thickness=4 mm, axial slices=44, flip angle=82°, field of view=240 x 240 mm). Anatomical images were acquired using a T1-weighted 3D gradient echo sequence (TR=2,300 ms, TE=2.36 ms, slice thickness=1 mm, number of slices=176, flip angle=9°, field of view=250 x 250 mm). The fMRI data were converted to BIDS format and preprocessed using the standard fMRIPrep pipeline (version 20.2.0). Preprocessing included surface reconstruction, spatial normalization to the MNI152NLin2009cAsym template, co-registration, and slice timing correction. Susceptibility distortion correction was not applied. Confound variables such as framewise displacement (FD), DVARS, and CompCor components were estimated. The EPIs from the fMRIPrep outputs were intensity normalized by applying each individual’s MNI-aligned brain mask. We then regressed out the following nuisance variables from all voxel time series within the brain mask: 24 head motion parameters, global signal, CSF, white matter, top-five cCompCor, top-five wCompCor, and linear trend. Frame censoring was applied using nearest-neighbor interpolation to replace frames with high motion (FD>0.5 mm, standardized DVARS>1.5, or frames identified by fMRIPrep as non-steady-state volumes). On average, 7.20% of frames across the entire experiment (including video watching, self-report, and resting) and 6.05% from the video-watching part were censored. Finally, spatial smoothing was applied using a 4 mm full-width at half maximum (FWHM) Gaussian kernel. Next, we selected TRs from the video-watching phases (15 phases in total across two runs). Accounting for the hemodynamic delay of 3-5 seconds (11, 46), we selected TRs from 4 seconds (2 TRs) after the start of each video to 4 seconds (2 TRs) after its end. The time-series data were than concatenated in a fixed order (randomly pre-determined, consistent across participants).

To ensure that participants remained awake during video-watching, we monitored them using a camera and excluded 18 videos from the analysis, in which 6 participants fell asleep. Additionally, we excluded videos with a mean framewise displacement (FD) exceeding 0.2 mm. If a participant had more than one-third of their videos excluded (i.e., five or more videos), that participant was excluded from the fMRI analysis, resulting in the removal of eight participants. Consequently, data from 53 participants were included in the final fMRI analysis, with 18 specific video blocks excluded across 6 of these remaining participants.

**Behavioral Data Analysis**

To examine the within-subject relationship between craving and self-relatedness, Pearson correlation coefficients were computed for each participant across the 15 video trials. These individual correlation coefficients were Fisher z-transformed prior to averaging, and a two-tailed t-test was performed to assess whether the mean correlation coefficient was significantly different from zero. The reported mean within-subject *r* reflects the back-transformed group-level z mean. To test the relationship between alcohol use severity and craving responses, we calculated the mean value for each participant and performed the Pearson correlation with the severity score. The same procedure was applied to examine the relationship between alcohol use severity and self-relatedness responses.

Then, to compare the explanatory power of self-relatedness and alcohol use severity in accounting for craving, a standardized multiple linear mixed-effects model was fit with craving as the dependent variable. The model included self-relatedness, alcohol use severity, and their interaction as fixed effects. The random effects included random slopes and intercepts for self-relatedness by participant to account for individual variability, as each participant provided 15 self-relatedness responses. Alcohol use severity was not modeled with random effects due to its single value per participant.

The mediation model was defined with alcohol use severity as the independent variable (X), mean craving responses as the dependent variable (Y), and mean self-relatedness responses for each participant as the mediator (M). Using the mediation package in R, we first fit a linear regression model to assess the effect of X on M ($M \sim a\cdot X$). Next, a second regression model was fit to evaluate the direct effects of both X and M on Y ($Y \sim b\cdot X+c\cdot M$). The mediation effect was calculated as the product of coefficients $a$ and $c$ from the two models ($a\cdot c$). The coefficient and the significance of the mediation effect was estimated based on 10,000 bootstrapped samples.

**Permutation Testing for IS-RSA**

To test the significance of inter-subject representational similarity analysis (IS-RSA) values, we conducted permutation testing with 10,000 permutations. For each permutation, we randomly shuffled the subject labels of the behavioral data, re-created the behavioral similarity matrix with the shuffled data, and recalculated the correlation with the neural similarity matrix, generating a null distribution of 10,000 IS-RSA values (Spearman rho-values) for each neural similarity matrix. Using this null distribution, we calculated a two-tailed p-value for each IS-RSA value. We applied Benjamini-Yekutieli (BY) correction (1) to adjust for multiple comparison across the 268 ROIs.

**Multiple linear regression analyses using similarity matrices**

To evaluate whether the identified relationships remained significant after controlling for alternative behavioral similarity measures, we conducted multiple linear regression analyses. In this model, the dependent variable was the neurobiological craving signature (NCS) similarity matrix, and the independent variables included both the primary behavioral similarity matrix—which previously demonstrated a significant correlation in the IS-RSA—and the behavioral similarity matrices used as controls. An interaction term between the independent variables was initially included but subsequently removed if found to be non-significant (2,3). To fit the regression model, we reshaped each similarity matrix into a vector of pairwise similarity values, yielding 1,378 values (i.e., _53_C_2_). Significance of the beta coefficients was assessed by permutation testing with 10,000 permutations. For each permutation, we randomly shuffled the subject labels of the neural data, re-created the pairwise neural similarity values, and re-ran the regression model to generate null beta coefficient values. This process yielded a null distribution of 10,000 permuted coefficients for each beta coefficient, from which we estimated a non-parametric p-value of the corresponding beta coefficient.

**Mediation analyses using similarity matrices**

To better understand how shared drinking motives influence neural synchrony and the role of craving responses, we tested whether NCS synchrony mediated the relationship between drinking motives and craving. In this model, the independent variable (X) was defined as the behavioral similarity matrix that demonstrated a significant result in the IS-RSA using the NCS (i.e., similarity matrix of drinking motives and self-relatedness, respectively). The mediator (M) was the NCS synchrony during video-watching, and the dependent variable (Y) was the similarity in craving responses. To fit the model, we reshaped each similarity matrix into a vector of pairwise similarity values, yielding 1,378 values (53 combination 2). Using the mediation package in R, we first fit a linear regression model to assess the effect of X on M ($M \sim a\cdot X$). Next, a second regression model was fit to evaluate the direct effects of both X and M on Y ($Y \sim b\cdot X+c\cdot M$). The mediation effect was calculated as the product of coefficients $a$ and $c$ from the two models ($a\cdot c$). The coefficient and the significance of the mediation effect was estimated based on 10,000 bootstrapped samples.

**Supplement 1. Validation of the Video-Watching Paradigm**

To validate that alcohol-drinking videos induce craving levels comparable to alcohol images, we collected behavioral responses from a separate set of participants. Twenty-two individuals from Seoul National University or surrounding community participated in a behavioral experiment conducted at a laboratory (14 females, age range: 19 to 46, mean age 25.41; all native Korean speakers). All participants were adults who self-identified as having problematic alcohol use (e.g., binge drinking or blackouts) but were not necessarily diagnosed with Alcohol Use Disorder (AUD). Based on the Alcohol Use Disorders Identification Test (AUDIT; (4)), 14 participants met the criteria for AUD (AUDIT score>10), six were classified as hazardous drinkers (AUDIT score>3), and two did not meet the threshold for risky drinking. Consequently, 91% of the participants exhibited risky drinking behavior based on the AUDIT. None of the participants had severe mental disorders, including substance use disorders, except for alcohol use disorder. All participants provided written informed consent and were compensated for their participation. The study was conducted in accordance with the Institutional Review Board at Seoul National University (IRB No. 2302/002-002).

Participants watched sixteen different videos featuring naturalistic alcohol consumption scenes (23.900 minutes in total, 69 to 106 seconds, mean=89.625, SD=9.824) and viewed sixteen images of alcoholic beverages for 7 seconds each (***Figure S1A***). Each video showed different characters drinking alcoholic beverages in various contexts. Stimuli were selected considering several features: social context (8 social drinking, and 8 solo drinking videos), gender of characters (8 male-only, 6 female-only, and 2 mixed-gender videos), type of alcohol (2 soju, 9 beer, 3 wine, 2 makgeolli-drinking videos), and food presence (11 videos with food, and 5 without food). All videos were extracted from Korean observational variety shows, 11 videos from “*I Live Alone*” and 5 videos from “*My Little Old Boy*.” Each image corresponded to the exact alcoholic beverage featured in its respective video. The order of videos and images was randomized. Participants either watched all 16 videos first, followed by the 16 images, or vice versa, as the order was counter balanced.

After watching each video, participants answered nine questions related to craving, self-relatedness, immersion (or concentration), boredom, perceived emotional valence of drinking reasons in the video (i.e., whether the drinking reasons portrayed in the video are positive or negative), the video’s naturalness, preference for the alcoholic beverage and the foods featured in the video, and whether they had seen the video before (***Figure S1A***). After viewing each image, they answered four questions regarding craving, immersion (or concentration), boredom, and preference for the alcoholic beverage. These questions are similar to those asked for the videos; only these four applied to images, as the other five questions are specific to videos that contain contexts. The questions were always presented in the same fixed order.

As a first step in validating our use of video stimuli, we compared subjective responses to four common questions across both video-watching and image-viewing paradigms: craving, immersion, boredom, and preference for alcoholic beverages. To examine whether stimulus type (video vs. image) had a significant effect on the responses, we fitted four standardized linear mixed-effects models. In each model, the dependent variable was each response variable, stimulus type was the fixed effect, and stimulus item and participant variables were included as random effects. Participants reported greater craving (*β*=0.258, *t*=4.079*, P*<0.001), immersion (*β*=0.762, *t*=12.889*, P*<0.001), and preference for the presented alcoholic beverages (*β*=0.329, *t*=5.044*, P*<0.001), as well as less boredom (*β*=-0.808, *t*=-13.430*, P*<0.001) after watching videos compared to viewing images (***Figure S1B***). Since the image of alcohol corresponded to the alcoholic beverage presented in the video, these results suggest that video stimuli induced greater craving and preference for alcoholic beverages while increasing engagement and reducing boredom.

Next, we investigated potential order effects on the response variables to determine whether responses were influenced by the order in which stimuli were presented. We ran nine standardized linear mixed-effects models, with each response variable as the dependent variable, the order of the videos as the fixed effect, and participant variable as a random effect. A response variable was considered affected by order if it was significantly explained by the order variable. All mixed-effects models were fit using the lmer function from the lmerTest package in R, and we applied Bonferroni correction to adjust for multiple comparison across response variables. As videos are presented later in the order, immersion decreased (*β*=-0.165, *t*=-3.934*,* corrected *P*=0.001) and boredom increased (*β*=0.137, *t*=2.924*,* corrected *P*=0.033). However, order had no effect on craving (*β*=-0.036, *t*=-0.741, corrected *P*=1.000), self-relatedness (*β*=−0.007, *t*=-0.137, corrected *P*=1.000), valence of drinking reasons (*β*=-0.076, *t*=-1.463, corrected *P*=1.000), naturality (*β*=-0.047, *t*=-0.906, corrected *P*=1.000), preference for alcoholic beverages (*β*=0.108, *t*=2.23, corrected *P*=0.238), preference for food (*β*=0.010, *t=*0.194, corrected *P*=1.000), and familiarity (*β*=-0.059, *t*=-1.13, corrected *P*=1.000). These results suggest that, except for the decrease in engagement over time, the order of presentation did not significantly influence participants' other responses.

Additionally, we examined whether the duration of each stimulus, defined as the time it was presented to participants, influenced the response variables. We ran another nine standardized linear mixed-effects models, with each response variable as the dependent variable, the duration of each stimulus as the fixed effect, and participant variable as a random effect. A response variable was deemed to be affected by duration if it was significantly explained by the duration variable. The results showed no significant effects of video duration on any of the response variables. Specifically, duration had no effect on craving (*β*=0.081, *t*=1.655, corrected *P*=0.889), self-relatedness (*β*=-0.012, *t*=-0.225, corrected *P*=1.000), concentration (*β*=0.038, *t*=0.897, corrected *P*=1.000), boredom (*β*=-0.007, *t*=-0.146, corrected *P*=1.000), valence of drinking reasons (*β*=-0.042, *t*=-0.814, corrected *P*=1.000), naturality (*β*=-0.045, *t*=-0.854, corrected *P*=1.000), preference for alcoholic beverages (*β*=0.042, *t*=0.862, corrected *P*=1.000), preference for food (*β*=0.024, *t*=0.449, corrected *P*=1.000), and familiarity (*β*=-0.014, *t*=-0.275, corrected *P*=1.000).

We also tested familiarity effects to assess whether participants' responses differed based on whether they have seen the video prior to the experiment. We fit eight standardized linear mixed-effects models, with each response variable (excluding the familiarity variable) as the dependent variable, familiarity responses as the fixed effect, and participant variable as a random effect. A response variable was considered to be influenced by familiarity if it was significantly explained by the familiarity variable. Only a small number of participants reported having seen any of the full videos before. One participant recognized two, and three others recognized one (***Figure S2A***). None of the response measures showed significant familiarity effects (***Figure S2B***): craving (*β*=0.110, *t=*2.164, corrected *P*=0.249), self-relatedness (*β*=0.046, *t*=0.867, corrected *P*=1.000), immersion (*β*=0.031, *t*=0.697, corrected *P*=1.000), boredom (*β*=-0.113, *t=*-2.307, corrected *P*=0.173), valence of drinking reasons (*β*=0.136, *t*=2.566, corrected *P*=0.086), naturality (*β*=0.018, *t*=0.330, corrected *P*=1.000), preference for alcohol (*β*=0.063, *t*=1.249, corrected *P=*1.000), and preference for food (*β*=0.076, *t*=1.427, corrected *P*=1.000). These findings indicate that participants' familiarity with the videos did not significantly alter their responses.

Lastly, we analyzed the associations among all response variables from the video-watching paradigm to identify potential confounding factors (***Figure S3***), focusing on their influence on craving. To estimate Pearson correlations between the eight response variables, we first computed the correlations for each individual, as each participant provided sixteen responses, one for each video. Higher craving was associated with higher self-relatedness (*r*=0.503, corrected *P*<0.001), greater immersion (*r*=0.369, corrected *P*<0.001), lower boredom (*r*=-0.4806, corrected *P*<0.001), positive reasons of the characters drinking alcohol (*r*=0.517, corrected *P*<0.001), higher naturality (*r*=0.382, corrected *P*<0.001), greater preference for alcohol (*r*=0.539, corrected *P*<0.001), and preference for food (*r*=0.405, corrected *P*<0.001). Greater immersion was strongly linked to lower boredom (r=-0.74, corrected *P*<0.001), and boredom was negatively correlated with all other response variables. In addition, preferences for alcohol and food were moderately correlated with each other (r=0.185, corrected *P*<0.001), indicating that participants with a higher preference for one type of consumption tended to rate the other positively.

In summary, these findings validate the use of naturalistic video cues for studying cue-induced alcohol craving. The videos induced sufficient levels of craving, with no unexpected confounding factors affecting craving levels. Among the sixteen videos, one with the lowest craving level was excluded from the fMRI paradigm due to time constraints.

**Supplement 2. Processing Speech on Alcohol Drinking Motives**

**Speech Data Processing**

Recorded speech from the free speech task (***Figure 1A***) was transcribed into text. The transcriptions were manually cleaned by removing starting remarks (i.e., “The reasons for drinking alcohol are...”) and concluding remarks (i.e., “In conclusion,” “These are the reasons that I drink alcohol.”), which appeared in most participants' responses. We next tokenized the text (with stop words removed (1)) and created embedding vectors. To select the embedding model that best captures individual differences, we compared multiple combinations of tokenization and embedding models. We evaluated three tokenization models: Open Korean Text (Okt)^[1]^, Korean morpheme analyzer (KKma)^[2]^, and Byte Pair Encoding (BPE)^[3]^. For embedding models, we tested three models: Sentence Transformer^[4]^, Korean Bidirectional Encoder representations from Transformers (KoBERT)^[5]^, and Korean Bidirectional Encoder representations from Transformers (KorBERT)^[6]^. The Sentence Transformer is a BERT-based model with up to 110 million parameters, comprising 12 layers, 768 hidden units, and 12 attention heads. It is pre-trained on large text corpora, including Wikipedia and BookCorpus. KoBERT has 92 million parameters, 12 layers, 768 hidden units, a feed-forward hidden size of 3,078, and 12 attention heads. It was trained on Korean Wikipedia, which contains approximately 5 million sentences and 54 million words. Similarly, KorBERT has 92 million parameters, 12 layers, a hidden size of 768, and 12 attention heads. It was trained on Korean Wikipedia and news datasets. This resulted in nine combinations of tokenization and embedding models (3 x 3).

**Model Comparison and Model Selection**

To determine the best-performing model, we treated human-rated speech data as the ground truth. While embedding vectors are known to effectively capture semantic structure (5), we aimed to validate whether they also reflect individual-level differences–particularly those perceptible to human raters. Three authors (M. K., S. S., and H. L.) independently rated each participant's speech based on nine predefined criteria for drinking motives: [1] drinking due to social pressure, [2] drinking socially without social pressure, [3] drinking alone, [4] drinking because of food, [5] drinking with food (but not necessarily because of it), [6] drinking due to negative emotion, [7] drinking due to positive emotion, [8] drinking out of habit, and [9] drinking for the taste of alcohol. Ratings were binary, and multiple criteria could apply to each participant. For instance, if a participant mentioned both drinking with friends and drinking alone, both “drinking socially without social pressure” and “drinking alone” were marked as 1. Raters reached consensus through discussion after independent coding.

We constructed a similarity matrix based on the human ratings by calculating Jaccard similarity between the 9-dimensional ratings for all participant pairs. We also created similarity matrices for each of the nine embedding vectors using cosine similarity between the embedding vectors for all participant pairs. We then computed the Spearman correlations between similarity matrix of the human-rated speech and each speech embedding vector (***Figure S4A***). Based on these comparisons, the combination of BPE tokenization and sentence transformer showed the highest correlation with human-ratings (***Figure S4B***) and was therefore selected for the main analysis.

**Analysis using Human-rated Alcohol Drinking Motives**

Although we selected the embedding model that best captured individual differences as perceived by human raters, the results derived from embedding vectors and human ratings may still diverge. The embedding vectors were 768-dimensional, while the human ratings consisted of binary responses across 9 dimensions. This discrepancy reflects two distinct approaches: embedding vectors capture fine-grained, individual-specific variation without relying on predefined constructs, whereas human ratings are interpretable but limited to preset criteria. Therefore, the choice between them should be guided by the research question—whether the goal is to detect subtle individual-level differences or to prioritize interpretability. As our primary aim was to capture unique individual variation in speech, we used embedding vectors in the main analyses. However, for comparison, we repeated all analyses using human-rating speech data: whole-brain IS-RSA (***Figure 2C***), IS-RSA using the NCS (***Figure 2E***), mediation analyses (***Figure S5-S6***), and multiple linear regression models predicting dynamic ISC (***Figure 4***).

Corresponding to “***Synchronized Brain Regions during Video-watching When Alcohol Drinking Motives are Similar Across Individuals****”*, we conducted IS-RSA to identify brain regions where activations are synchronized across individuals with more similar human-rated drinking motives. A behavioral similarity matrix was created by computing Jaccard similarities across participants’ binary human-rated motive profiles. Neural similarity matrices were computed for each of the 268 regions of interests (ROIs) based on Shen et al.’s (6) brain parcellation. We then computed the Spearman correlation between the behavioral similarity matrix and the neural similarity matrices across all ROIs. Although both speech embedding vector and human ratings were derived from the same speech, unlike embedding vectors (***Figure 2C***), similarity in human-rated speech motives did not predict neural synchrony in any brain region.

Similarly, in the IS-RSA using the NCS (corresponding to “***The Neurobiological Craving Signature (NCS) is synchronized during Video-watching When Alcohol Drinking Motives are Similar****”***)**, similar human-rated motives did not predict NCS synchrony during video-watching ($\rho$=-0.005, *P=*0.928). Given that human-rated speech did not explain NCS activity, we did not conduct further mediation analyses using these human ratings.

Finally, we conducted an analysis corresponding to “***Diverging Drinking Motives Lead to Idiosyncrasy in the NCS in Response to Alcohol Cues****”*. To further examine individual differences in the NCS and test whether human-rated motives could explain such difference, we used the same participant-by-participant beta matrices for alcohol and food (derived from multiple linear regression models; ***Figure 4A***), which represent the extent to which neural synchrony is explained by cue presence (***Figure 4B***). To assess whether similarity in human-rated motives could explain these cue-related beta coefficients, we computed Spearman correlations between each beta matrix and the human-rated motives similarity matrix. The results mirrored those from the main analysis using motives represented by embedding vectors (***Figure 4C***): similarity in $\beta_{alcohol}$ was significantly associated with human-rated motives ($\rho$=0.067, *P=*0.047), whereas $\beta_{food}$ showed no such relationship ($\rho$=0.024, *P=*0.469). Moreover, absolute beta values showed a consistent pattern across both types of speech-derived measures (***Figure 4D***). The most dissimilar pairs showed significant low $\beta_{alcohol}$ (mean of $\beta_{alcohol}$ in the first quantile=-0.019, 95% CI=[-0.036, -0.002], *t*=-2.182, *P=*0.030). These findings suggest that in subject pairs with the most divergent drinking reasons, NCS responses to alcohol cues diverge (i.e., desynchronize). Notably, this pattern was observed regardless of whether drinking motives were represented by embedding vectors or human ratings.

In sum, while human ratings offered interpretable representations of alcohol drinking motives, they did not account for neural synchrony at the whole-brain level or NCS synchrony during naturalistic video-watching. They did, however, capture individual variability in cue-related NCS synchrony—unlike other behavioral measures such as craving, self-relatedness, alcohol use severity, or AUD symptom profiles—suggesting that human ratings do capture certain aspects of individual variability in responses to alcohol-related cues. In contrast, speech embedding vectors consistently revealed significant associations across analyses, suggesting that embedding vectors captured unique, fine-grained variance in neural responses linked to drinking motives—variance that was not accessible through human ratings, despite being based on the same speech data. These findings highlight the utility of embedding-based approaches in detecting subtle individual differences, particularly when the research goal is to uncover idiosyncratic patterns across individuals.

**Supplement 3. Analysis using Nearest-Neighbor model to compute similarity in alcohol use severity**

To compute similarity across individuals in terms of alcohol use severity, we applied the Anna-Karenina (Anna-K) model, which defines similarity as the average of two participants’ severity ranks, normalized by the total number of participants. Specifically, similarity between participant *i* and participant *j* was calculated as:

$${Similarity}_{AnnaK}= {(Rank}_{i}+ {Rank}_{j})\div2 \div N$$

where ${Rank}_{i}$ and ${Rank}_{j}$ are the severity ranks of the two participants (severity scores were rank-transformed across participants, with Rank=1 indicating the lowest severity and Rank=N indicating the highest), and $N$ is the total number of participants (here, $N$=53). This model is inspired by the Tolstoy's line, “All happy families are alike; each unhappy family is unhappy in its own way.” According to this model, individuals with higher severity scores are assigned higher similarity values, while those with lower severity scores are assigned lower similarity values. On the other hand, the Nearest Neighbor (NN) model computes similarity based on the absolute difference between ranks:

$${Similarity}_{NN}=1- \left| {Rank}_{i}- {Rank}_{j} \right|$$

where ${Rank}_{i}$ and ${Rank}_{j}$ are defined identically as in the Anna-K model. The NN model assumes that individuals are most similar to their nearest neighbors in severity, regardless of their absolute severity levels. Thus, individuals with similar ranks, whether high or low, receive higher similarity scores.

In the main analyses, we used the Anna-K model to assess similarity in alcohol use severity, as our primary interest was whether higher severity could explain cue-induced craving, as suggested by prior literature (7–11). Here, as a supplementary analysis, we repeated the same set of analyses using the NN model. In the main text, none of the findings using the Anna-K model were statistically significant, and consistent with this, no significant findings emerged using the NN model. Specifically, no brain regions showed synchronized activation based on NN-based similarity in alcohol use severity (*corresponding to “****Synchronized Brain Regions during Video-watching When Alcohol Drinking Motives are Similar Across Individuals****”*), and NCS synchrony was not associated with NN-based alcohol use severity ($\rho$=0.026, *P=*0.280; *corresponding to “****The Neurobiological Craving Signature (NCS) is synchronized during Video-watching When Alcohol Drinking Motives are Similar****”*). Given this lack of association, we did not proceed with mediation analyses using this variable. Finally, and consistent with the Anna-K results, NN-based alcohol use severity did not explain individual differences in NCS synchrony to alcohol cues ($\beta_{alcohol}$: $\rho$=0.006, *P=*0.811; $\beta_{food}$: $\rho$=-0.041, *P=*0.122; *corresponding to “****Diverging Drinking Motives Lead to Idiosyncrasy in the NCS in Response to Alcohol Cues****”*).

**Supplement 4. Validation of the Neurobiological Craving Signature (NCS)**

To validate the NCS in our dataset, we first fit a general linear model (GLM) using two main regressors: presence (i.e., onsets and durations) of alcohol cues and food cues in the videos (***Figure S7***). Additional regressors included onsets and durations of other screens presented (i.e., fixation, instruction, craving and self-relatedness responses), low-level visual and auditory features, and six motion regressors. We performed the first-level GLM analysis using a contrast “alcohol” and “food,” and computed a single NCS-weighted beta value per participant for each contrast (***Figure S8A***). Specifically, by taking the dot product of each voxel-wise beta map with the NCS, we obtained a single response score of the NCS for “alcohol” and “food” contrasts for each participant. We then compared the NCS activation for “alcohol” and “food” using a two-sampled t-test.

Next, we examined whether NCS synchrony (i.e., ISC of the NCS) was more correlated with the time series of alcohol cue presence than with that of food cue presence. To analyze this, we computed dynamic ISC (12) of the NCS time series using a sliding window method (***Figure S8C***). In this approach, the Fisher's z-transformed Pearson's correlation of pairwise participants' brain responses was computed within each window (window size=10 TR). This process was repeated across the entire video-watching duration by sliding the window one step at a time (step size=1 TR), resulting in a time series of dynamic ISC for every pair of subjects. We then averaged the dynamic ISC across pairs to obtain a single time series representing neural synchrony of the NCS during video watching (***Figure S8C***). To match the temporal dimension of the dynamic ISC, we applied the sliding window to the time-series of alcohol and food cues (the averaged ratings from 7 raters; ***Figure S7***) within each time window. Then, after standardizing each time-series data, we computed the Pearson correlation between the averaged dynamic ISC and time series of the alcohol cues, as well as between the dynamic ISC and the time series of the food cues (***Figure S8D***). The significance of these correlations was tested using a non-parametric circular shift randomization method. Given the specific constraints of the time series length (N=700 TR), we conducted an exhaustive permutation test by calculating correlation coefficients for all possible N unique circular shifts. For each unique shift (From 0 to N-1), the dynamic ISC time series was circularly shifted by that amount to generate a null distribution of correlation coefficients. Correlation coefficients were computed between the shifted time series and the unchanged cue time series to generate a null distribution for each cue. The p-value was estimated as the proportion of permuted correlation coefficients equal to or more extreme than the observed correlation coefficient.

The results revealed that NCS activation was higher during the presentation of alcohol cues compared to food cues by fitting the GLM (*t=*3.560, *P*<0.001; ***Figure S8B***; see ***Figure S9*** for the whole-brain second-level results). Next, by conducting correlation analyses using the time-series of NCS synchrony (i.e., dynamic ISC) and the presence of cues during video-watching (***Figure S8C***), we found that the NCS was more synchronized in the presence of alcohol (*r=*0.190; *P=*0.026), and less synchronized in the presence of food (*r=-*0.251; *P=*0.005; ***Figure S8D***). Supporting the specificity of the NCS, these patterns were not replicated with other alternative neuromarkers, the details of which are described in ***Supplement 6***.

**Supplement 5. Additional Mediation Analyses using Neural synchrony of Specific Conditions during Video-watching**

Given that each video combined alcohol cues, food cues, and neutral contextual cues, we investigated whether the presence of specific alcohol cues accounted for the mediation effect (***Figure S5***) by isolating specific cue-related timepoints from the NCS time-series. Seven independent raters evaluated whether alcohol-related and food-related cues appeared with motion in each video, assessing them in real-time while watching videos. We focused on cues with motion rather than static cues, as some cues are so trivial and not intended to draw attention. For the alcohol cues, the raters reached agreement on 75.31% of all time points (51.73% from no alcohol, 23.57% from yes alcohol). For the food cues, agreement was achieved on 86.42% of the time points (69.69% from no food, 16.73% from yes food). To standardize the data, we rounded the averaged ratings (***Figure S7A***) to binary values (0 or 1) at each time point (***Figure S7B***). For each condition, we annotated the relevant TRs from the videos and selected TRs that were 4 seconds (2 TRs) after cue onset to account for the hemodynamic delay. We then concatenated the TRs within each condition, following prior work (13,14), and extracted the corresponding NCS time series for each participant.

We separately analyzed the activation of the NCS for five conditions: alcohol cues, food cues, alcohol cues with contexts, food cues with contexts, and contexts. The durations for each condition were 548 seconds (274 TRs) for “alcohol cues” and 294 seconds (147 TRs) for “food cues.” Overlapping TRs were excluded from both conditions to isolate the distinct effects of alcohol and food cues. For “contexts,” TRs without the presence of any alcohol or food cues were selected, totaling 482 seconds (241 TRs). “Alcohol cues with contexts” included TRs without any food cues, totaling 1030 seconds (515 TRs), and “Food cues with contexts” included TRs without any alcohol cues, totaling 776 seconds (388 TRs). For each of the five conditions, we extracted the NCS time-series and repeated the mediation analyses with one modification: the mediator (M) was replaced by the NCS synchrony for each condition. Thus, we tested five additional mediation models. For example, the model for the “alcohol cues” condition examined whether similar behavior led to the NCS synchrony specifically during alcohol cue presentation and whether this synchrony corresponded to similar self-reported craving.

The mediation effect was significant only when the neural data included timepoints for alcohol cues—either alcohol cues alone (mediation effect=0.018, 95% CI=[0.001, 0.041], *P*=0.043; ***Figure S6A***) or alcohol cues with contexts (mediation effect=0.019, 95% CI=[0.003, 0.042], *P*=0.018; ***Figure S6B***)—and disappeared when timepoints for alcohol cues were removed (***Figure S6C-E***). Specifically, NCS synchrony in response to food cues was not a significant mediator of the relationship (mediation effect=0.005, 95% CI=[-0.001, 0.020], *P*=0.195; ***Figure S6C***). Similarly, NCS synchrony during the presence of food cues with contexts did not show a significant mediation effect (mediation effect=0.007, 95% CI=[-0.001, 0.023], *P*=0.129; ***Figure S6D***). The model incorporating NCS synchrony during the presence of contexts (without alcohol or food cues) as well, the mediation effect was not significant (mediation effect=0.002, 95% CI=[-0.002, 0.015], *P*=0.552; ***Figure S6E***). These findings suggest that neural synchrony specifically in response to alcohol cues was closely associated with similarities in alcohol drinking motives, which in turn contributed to similar self-reported craving after video-watching. To evaluate and compare model fit of the five models, we used Akaike Information Criterion (AIC) values (15), where lower AIC values indicate better fit. By using the difference in AIC values (ΔAIC), we assessed the relative performance of the models (***Figure S6F***), with an AIC difference greater than 10 indicating strong evidence for the model with the lower AIC (16). The mediation model incorporating NCS synchrony during the entire video-watching phase (***Figure S5A***) demonstrated the best fit with an AIC value of -11607.526. The next best-fitting model, which included alcohol cues with contexts, had an AIC of -11343.553. Compared to the model incorporating only alcohol cues (AIC=-10637.328), the AIC difference exceeded the threshold of 10, indicating that neural synchrony in response to contextual elements of alcohol drinking explains additional variance. However, it is noteworthy that only models including alcohol cues showed a significant mediation effect of NCS synchrony. These results underscore the importance of alcohol cues in mediating the relationship between drinking motives and craving responses.

**Supplement 6. Examining Specificity of the Relationship Between Speech Embedding Vectors on Alcohol Drinking Motives and the Neurobiological Craving Signature**

**Specificity of Speech Embedding Vectors on Alcohol Drinking Motives**

To examine the specificity of the speech embedding vectors derived from alcohol drinking motives, we repeated the identical analysis framework using two additional, duration-matched (3 minutes) speech samples collected from the same cohort of participants. These comparison samples comprised a speech about quitting drinking—elicited by the prompt “Please share your thoughts about quitting, refraining, or reducing drinking alcohol”—and a control speech elicited by the prompt “Please share what happened before you came to this laboratory today.” Following the exact preprocessing pipeline applied to the original speech on drinking motives (***Supplement 2***), we transcribed the speech into text, tokenized the text using BPE tokenization with stop word removal (17), and then generated embedding vectors using sentence transformer.

For each of these alternative speech data, we conducted IS-RSA using both the 268 whole brain ROIs and the NCS. While the original speech (i.e., speech on drinking motives) was associated with neural synchrony in the middle temporal gyrus, temporal pole, and posterior cerebellum during video-watching (***Figure 2C***), IS-RSA using 268 ROIs revealed no significant brain regions associated with either the quitting drinking or control speech embedding vectors. Similarly, while shared drinking motives explained NCS synchrony during video-watching (***Figure 2E***), IS-RSA using the NCS showed no significant results for either of the other speech types (quitting drinking: $\rho$=0.065, *P*=0.333; control speech: $\rho$=0.055, *P=*0.373). Notably, even after controlling for these speech embedding vectors individually, the embedding vector on drinking motives remained a significant predictor of NCS synchrony (***Table S3***). Mediation analyses were not conducted, as neither the quitting drinking nor control speech embedding vectors were significantly associated with the NCS. Lastly, while drinking motives explained individual differences in NCS synchrony in response to alcohol cues (***Figure 3***), neither the quitting drinking nor control speech embedding vectors explained the beta matrices for alcohol cues (quitting drinking: $\rho$=0.000, *P*=0.992; control speech: $\rho$=0.007, *P*=0.858) or food cues (quitting drinking: $\rho$=-0.012, *P*=0.748; control speech: $\rho$=0.010, *P=*0.787). Together, these findings support the specificity of the speech embedding vector on drinking motives in its relationship with the NCS, as its effects were distinct from those of the other speech types.

**Specificity of the Neurobiological Craving Signature**

To evaluate the specificity of the NCS, we repeated the entire analysis pipeline using three alternative neuromarkers: the Brain Reward Signature (BRS; (18)), the Picture-Induced Negative Emotion Signature (PINES; (19)), and the Neurologic Pain Signature (NPS; (20)). The PINES (<https://github.com/canlab/Neuroimaging_Pattern_Masks/tree/master/Multivariate_signature_patterns/2015_Chang_PLoSBiology_PINES>) and NPS (through email request to first author of (20)) consisted of unthresholded voxel weights, as the NCS, while the BRS (<https://github.com/canlab/Neuroimaging_Pattern_Masks/tree/962bc43af0b7dd4cba1a92d15eee09b1fc860a48/Multivariate_signature_patterns/2023_Speer_Brain_Reward_Signature_BRS>) consists of thresholded voxel weights. All neuromarkers used in this analysis were developed using LASSO-PCR with cross-validation, tested for sensitivity and specificity, and generalized to out-of-sample data. To match the value range with the NCS, we scaled the BRS voxel weights by multiplying them by 0.001 and the NPS voxel weights by multiplying them by 0.1, which does not alter the original distribution or interpretation of the voxels.

None of the findings observed using the NCS were replicated with these alternative neuromarkers, supporting the specificity of the NCS in its relationship with speech embedding vectors on drinking motives. First of all, unlike the NCS that exhibited preferential responsivity to alcohol cues compared to food cues (***Figure S8B & E***), none of the three alternative neuromarkers were more responsive to alcohol stimuli. Specifically, the BRS responded more strongly to food cues (*t*=-2.370, *P*=0.020; ***Figure S10A***), the PINES did not differentiate between alcohol and food cues (*t*=-0.008, *P*=0.994; ***Figure S10B***), and the NPS exhibited greater responses to food cues than alcohol cues (*t*=-9.073, *P*<0.001; ***Figure S10C***). Similarly, while the dynamic ISC of the NCS was significantly correlated with alcohol cue presence (***Figure S8E***), no such pattern was observed for the other neuromarkers (BRS: *r*=0.189, *P=*0.098; PINES: *r=*0.058, *P=*0.282; NPS: *r=*0.237, *P=*0.077; ***Figure S11***). For food cues, only the PINES showed a significant negative correlation (*r=*-0.235, *P=*0.032), with no significant results for the BRS (*r=*-0.228, *P=*0.077) or NPS (*r=*-0.152, *P=*0.161). These findings highlight the specificity of the NCS to alcohol-related cues, providing initial evidence that the NCS generalized to the current dataset and could be used as a useful proxy for neural craving.

In the IS-RSA, while the NCS was significantly associated with self-relatedness and with alcohol drinking motives (***Figure 2E***), no such association was found in the other neuromarkers. The BRS (***Table S4***) and the PINES (***Table S5***) showed no significant association with any behavioral measure. The NPS was associated with two behavioral similarities. First, individuals with similar craving responses for each video showed synchronized activation in the NPS ($\rho$=0.116, *P*=0.010). Second, individuals with similar alcohol use severity based on the nearest neighbor model (NN; $\rho$=0.052, *P*=0.042; ***Table S6***)—reflecting similarity in severity scores, not necessarily higher severity—also showed synchronized NPS activation. These results remained significant even after controlling for other behavioral measures (***Table S7***). Given that only the NPS yielded significant results, we conducted a mediation analysis using the NPS synchrony as a mediator, with NN-based similarity in alcohol use severity as the independent variable, and similarity in craving responses as the dependent variable. The analysis revealed a significant mediation effect of NPS synchrony (mediation effect=0.011, 95% CI=[0.002, 0.023], *P*=0.013; ***Figure S12A***), linking similar alcohol use severity to similar craving responses. Mediation analyses distinguishing between alcohol cues, food cues, and contexts revealed that NPS synchrony mediated the link between similar alcohol use severity and craving responses, particularly when food cues were embedded within broader contexts (***Figure S12B–F***).

In the regression analysis investigating individual differences, unlike the NCS—whose dynamic ISC in response to alcohol cues was associated with drinking motives (***Figure 3***)—none of the other neuromarkers were significantly related to alcohol drinking motives (BRS: $\rho$=0.032, *P*=0.401; PINES: $\rho$=-0.010, *P*=0.838; NPS: $\rho$=0.012, *P*=0.734; ***Figure S13A-C***). While the PINES and NPS showed no significant results, the BRS exhibited a notable pattern (***Figure S13A***): its neural synchrony in response to alcohol cues tended to increase among individuals with similar self-relatedness responses ($\rho$=0.082, *P*=0.021). Conversely, its neural synchrony in response to food cues tended to decrease (become more diverse) among individuals with similar craving responses ($\rho$=-0.074, *P*=0.026). As illustrated in ***Figure S13D***, this suggests that individuals with similar self-relatedness responses showed increased neural synchrony related to reward at the presence of alcohol. Taken together, the findings from the NCS were not replicated with the BRS, PINES, or NPS, supporting the specificity of the NCS in its relationship with drinking motives.

**Figures and Tables**

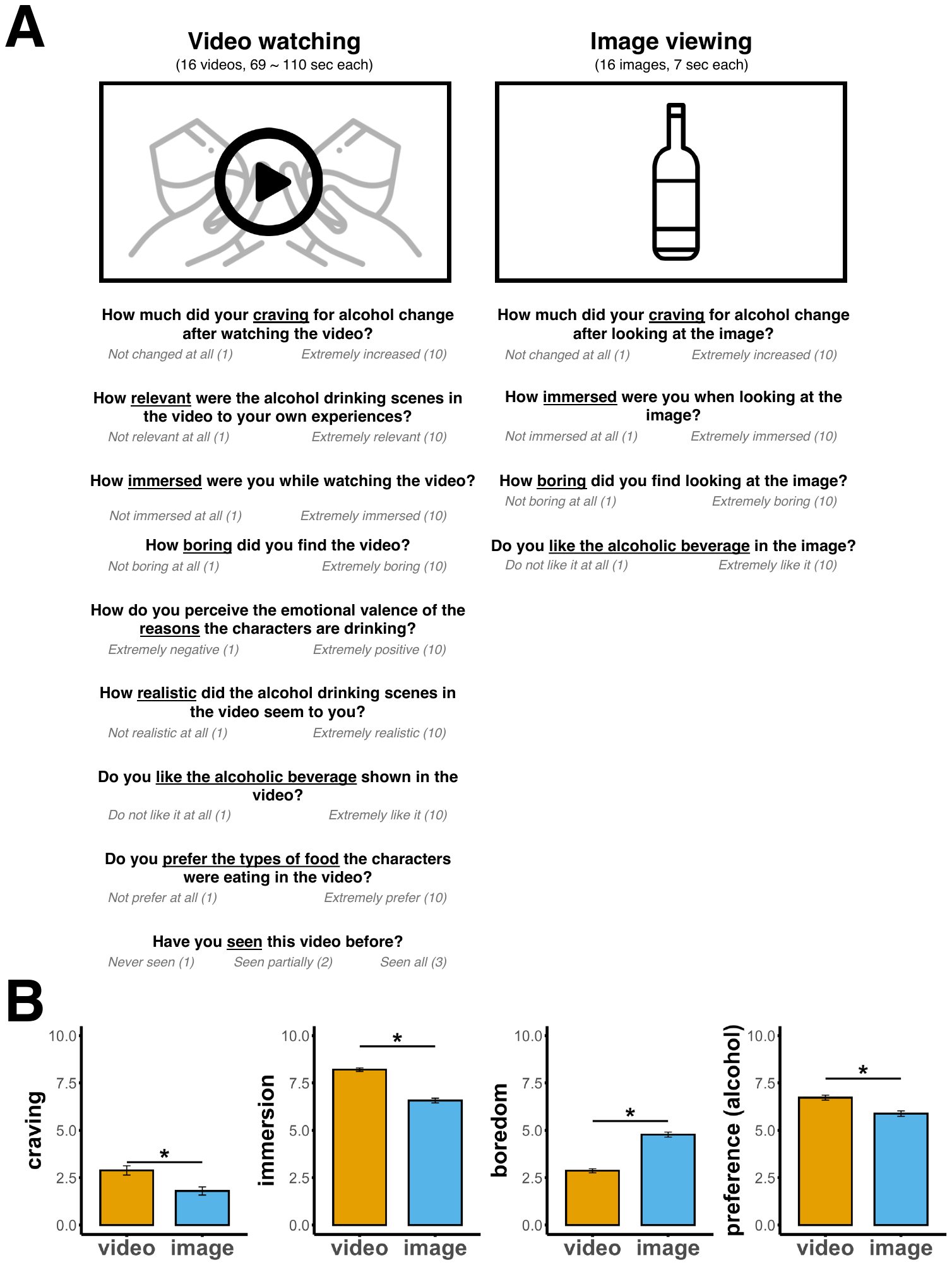

**Supplementary Figure S1.** Validation of naturalistic video-watching paradigm. *(A)* Video-watching and image-viewing paradigms. Participants watched sixteen videos, each lasting between 69 to 110 seconds, and answered nine questions after watching each video. Every video included scenes of alcohol consumption, and the order in which the videos were presented was randomized across participants. Participants also viewed sixteen images, each depicting the alcoholic beverage shown in the video for 7 seconds, and answered four questions. These questions were adapted from the video-watching paradigm and paraphrased to be applicable after viewing images. The order of the video-watching and image-viewing paradigms was counterbalanced across participants to prevent order effects. *(B)* Comparison of self-reported responses between video-watching and image-viewing paradigms, grouped by the same alcoholic beverage. Each comparison was based on a standardized linear mixed-effects model, with each response variable as the dependent variable and stimulus type (video vs. image) as the fixed effect, and stimulus item and participant variable as random effects. Asterisks indicate statistically significant differences between video and image (*P*<0.05, Bonferroni correction applied to adjust for multiple comparison across variables).

**Supplementary Figure S2.** Familiarity effects in the video-watching paradigms. *(A)* The figure presents participants' responses to the question, “Have you seen this video before?” to assess familiarity with each video. The X-axis represents each video item, and the Y-axis shows the number of responses for each option, with “never seen” represented in the darkest grey. *(B)* Familiarity effects were tested by fitting a standardized linear mixed-effects model for each response variable. The X-axis shows the level of familiarity, and the Y-axis displays the mean response for each variable. None of the responses showed a significant familiarity effect, as the familiarity variable did not significantly explain any of the eight variables.

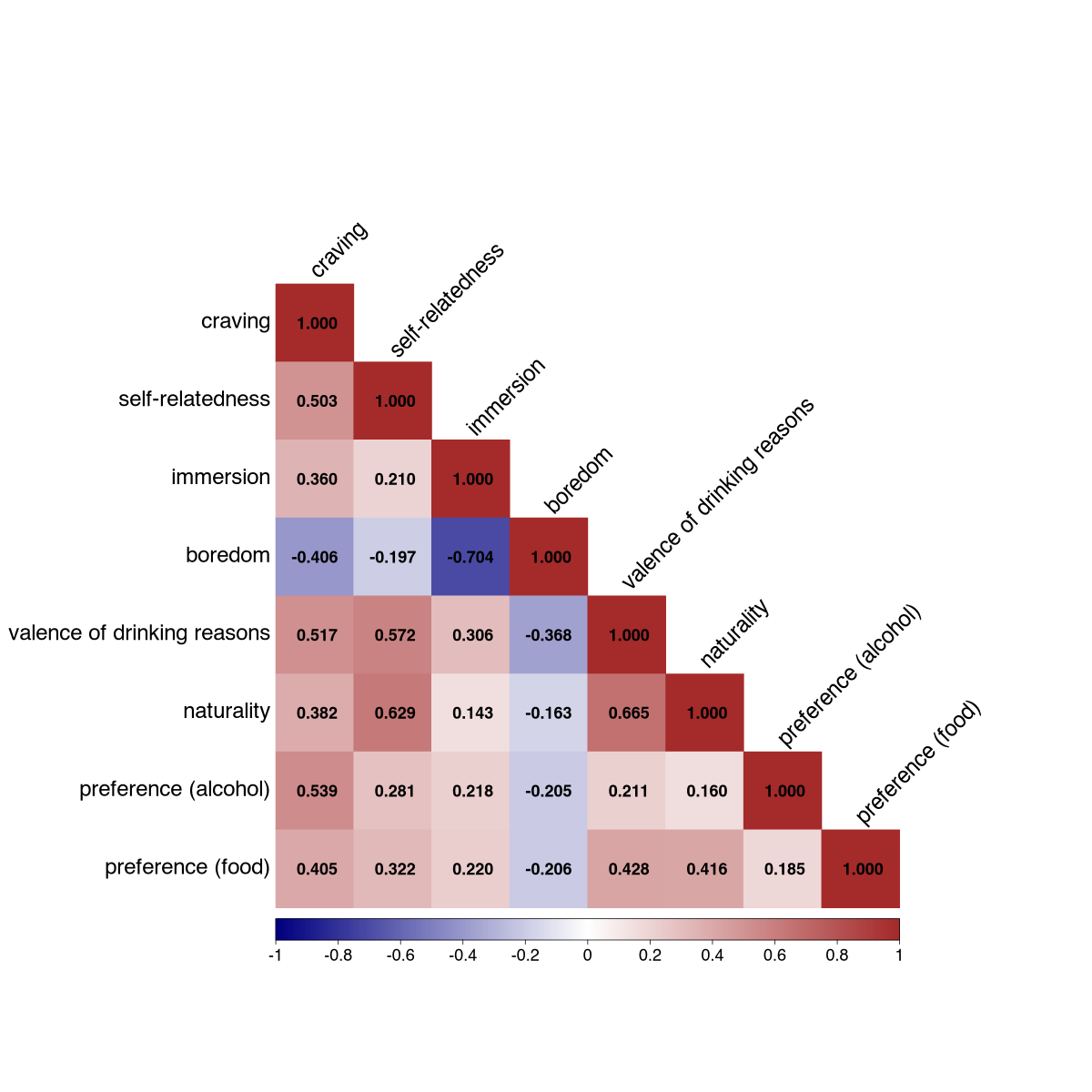

**Supplementary Figure S3.** Correlation matrix of response variables in the video-watching paradigm. Pearson correlation coefficients were first estimated for each individual, and the averaged coefficients across participants are presented in the figure. All correlation coefficients were significantly different from zero (*P*<0.05, Bonferroni-corrected to adjust for multiple comparisons across response variables).

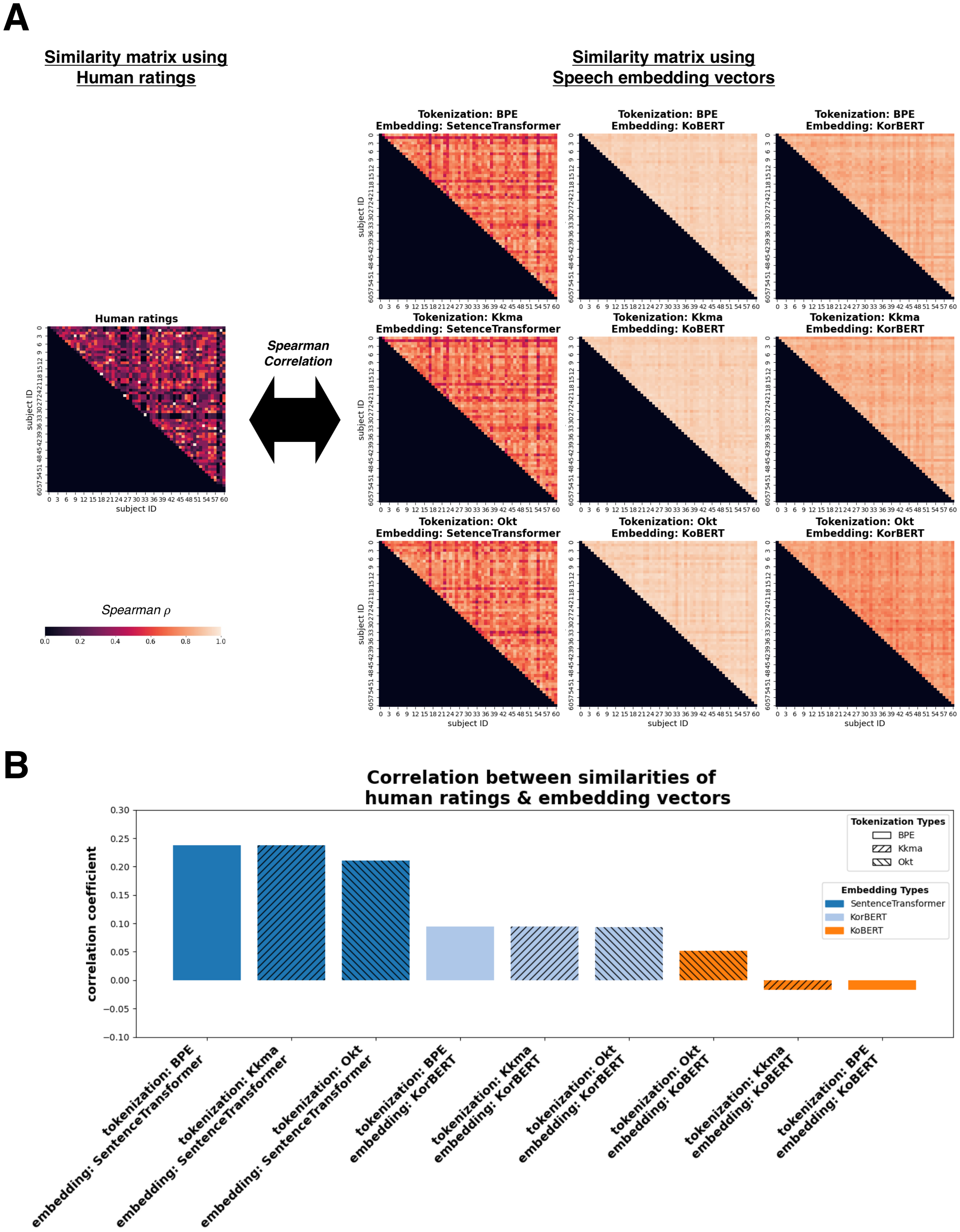

**Supplementary Figure S4.** Model comparison and model selection of embedding and tokenization models for speech data. *(A)* Comparison of the human-rated speech similarity matrix and speech embedding vector similarity matrices*.* To identify the best embedding vectors that represent individual differences, we compared human-rated speech on drinking reasons with the nine different embedding vectors of the transcribed speech (**Figure 1A**). The left matrix displays Jaccard similarity between human-rated speech across all participants, treated as the ground truth of individual differences. The right panel contains nine similarity matrices, each corresponding to a different combination of tokenization and embedding models. Tokenization methods tested include Byte Pair Encoding (BPE), Korean morpheme analyzer (KKma), and Open Korean Text (Okt). The embedding models tested include Sentence Transformer, KoBERT, and KorBERT. Cosine similarity was used to calculate similarity matrices for the embedding vectors. *(B)* Correlations between human-rated speech similarity matrix and speech embedding vector similarity matrices. Th*e* bar plot shows Spearman correlation coefficients ($\rho$) between the similarity matrix derived from human ratings and similarity matrices from different combinations of tokenization and embedding models (as shown in A). The X-axis lists each combination of tokenization and embedding model, and the Y-axis displays the corresponding correlation coefficients.

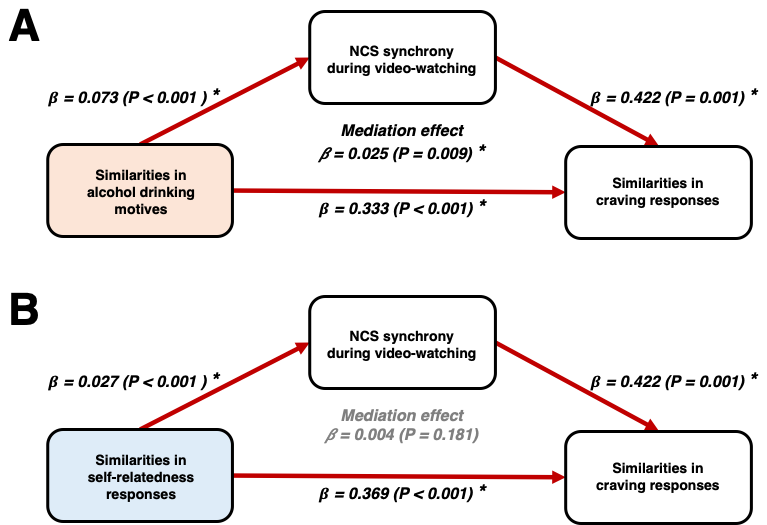

**Supplementary Figure S5.** Mediation analysis examining synchrony of the neurobiological craving signature (NCS) as a mediator between behavioral similarities and craving similarities. *(A)* Mediation analysis using similarities in alcohol drinking motives (speech embedding vectors) as the independent variable. *(B)* Mediation analysis using similarities in self-relatedness responses as the independent variable. The coefficient and significance of the mediation effect were assessed using bootstrapping with 10,000 samples. Asterisks indicate significant beta coefficients (*P<*0.05).

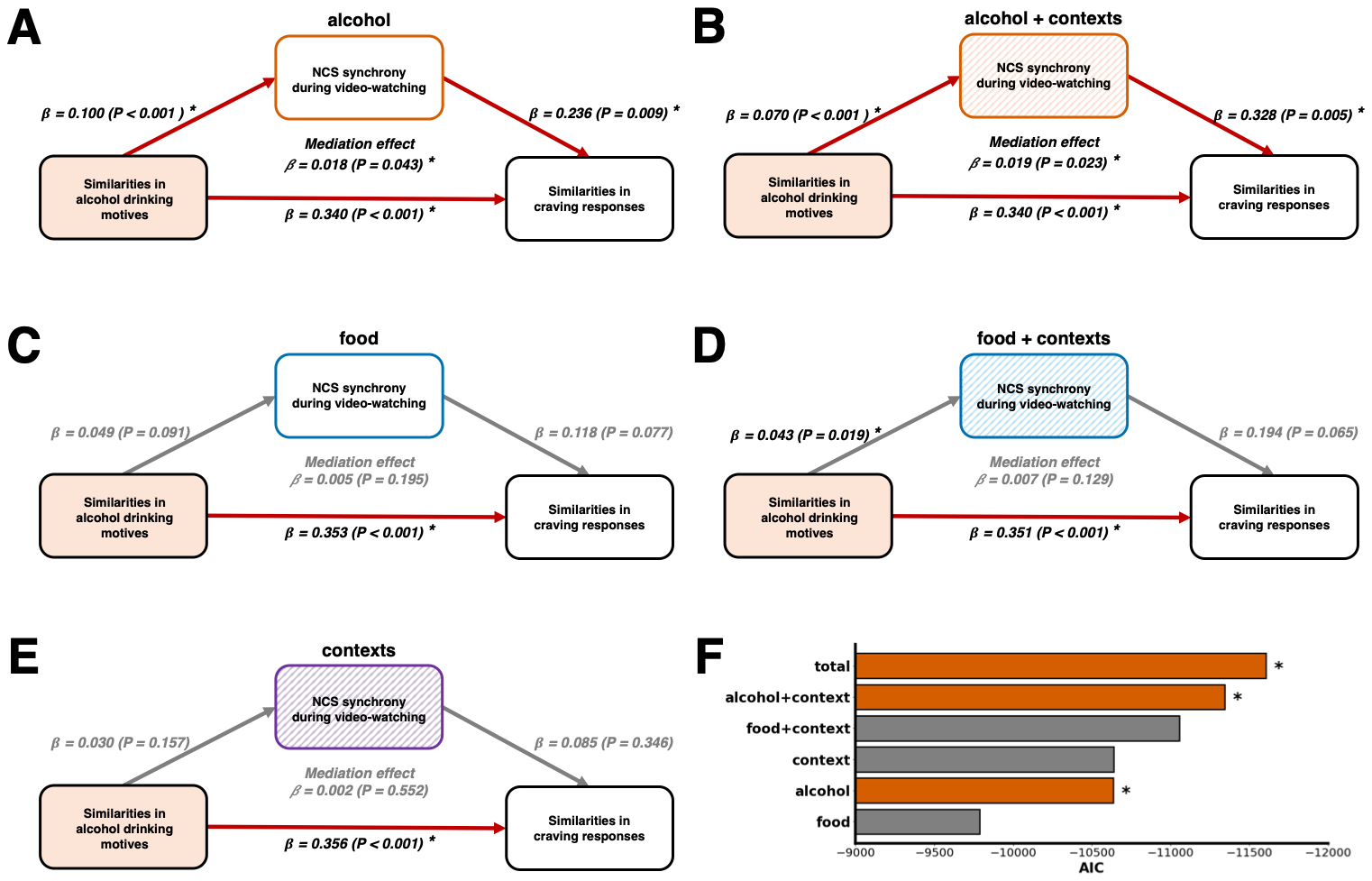

**Supplementary Figure S6.** Mediation analysis examining whether the relationship between similar alcohol drinking motives (speech embedding vectors) and similar craving responses is mediated by synchrony of the neurobiological craving signature (NCS) in response to *(A)* alcohol cues, *(B)* alcohol cues with contexts, *(C)* food cues, *(D)* food cues with contexts, and *(E)* contexts alone during video-watching. The coefficient and significance of the mediation effect were assessed using bootstrapping with 10,000 samples, with significant coefficients (*P<*0.05) indicated by asterisks. *(F)* Model comparison results for A, B, C, D, and E. The Y-axis shows the mediation models, with labels indicating the mediator in each model and specifying which information in the videos was included to assess the NCS synchrony. The X-axis displays the Akaike Information Criterion (AIC) values for each model, where lower AIC values indicate better model fit. Orange-colored bars denote models with significant mediation effects (*P<*0.05).

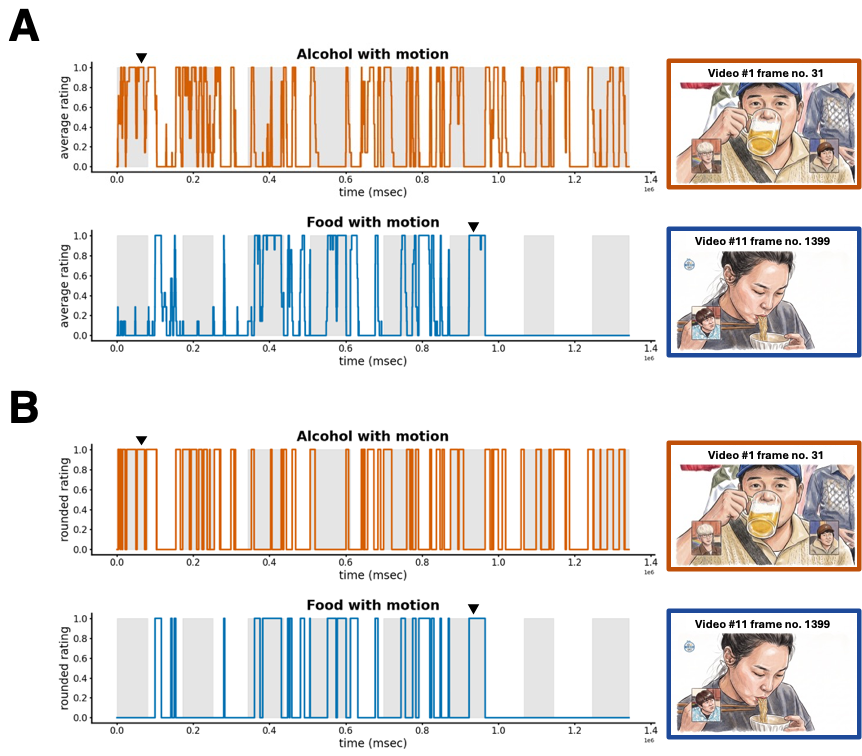

**Supplementary Figure S7.** Alcohol and food regressors extracted from the videos. *(A)* The line graphs show the presence of alcohol-related cues (alcohol with motion) and food-related cues (food with motion) over time, extracted from the 15 videos. Ratings from seven independent raters were averaged at each time point. The top graph (in red) displays the alcohol regressor, while the bottom graph (in blue) shows the food regressor. Gray and white background indicates change in the video items, ordered sequentially from the leftmost (Video #1) to the rightmost (Video #15). The images on the right are example computer-generated drawings substituted to illustrate video frames when alcohol (Video #1, frame 31) and food (Video #11, frame 1399) appeared with motion, as indicated by the black triangles on the line graphs. (*B*) The averaged ratings shown in (A) were rounded to binary values (0 or 1) at each time point, indicating presence (1) or absence (0) of alcohol or food cues with visible motion.

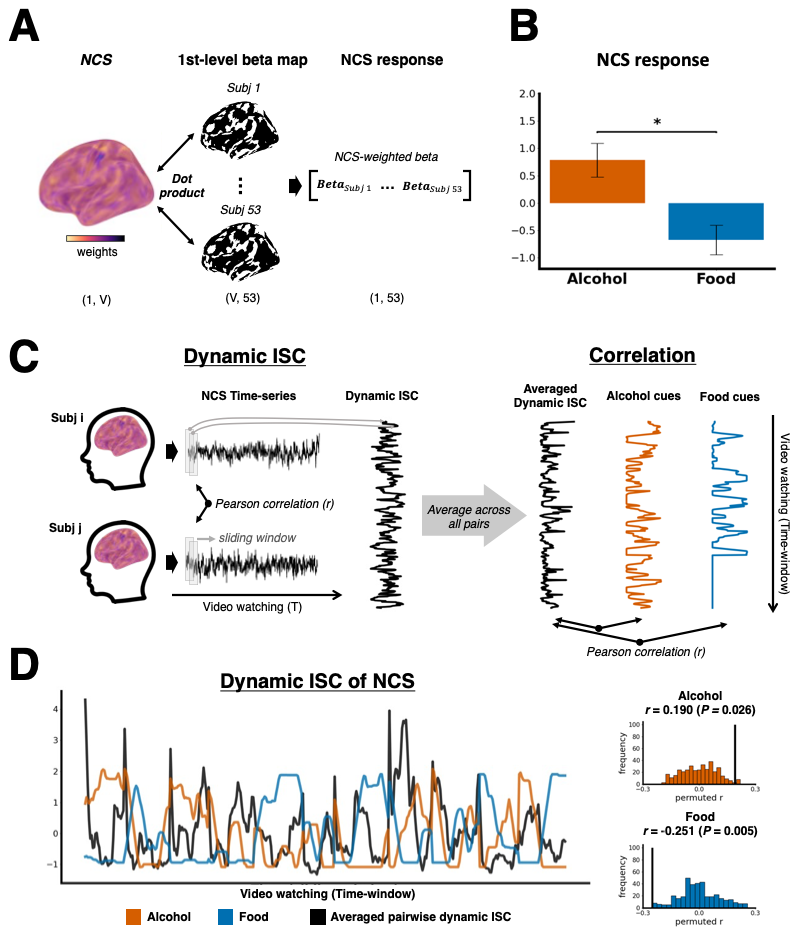

**Supplementary Figure S8.** Validation of the Neurobiological Craving Signature (NCS). *(A)* Computing NCS responses after fitting a first-level general linear model (GLM). For each participant, a single NCS-weighted beta value was computed by taking the dot product of the voxel-wise beta map from the GLM with the voxel-wise weights of the NCS. This value represents the NCS response to alcohol or food cues during video-watching. *(B)* Comparison of NCS responses to alcohol and food cues. After running the first-level GLM, a single NCS response was computed for each contrast (i.e., “alcohol” and “food”) per participant. The X-axis represents the first-level contrast of “alcohol” and “food,” and the Y-axis shows the mean NCS response across participants, with error bars indicating standard error. A two-sample t-test was performed to compare the NCS responses to alcohol and food cues. The asterisk indicates a significant difference (*P*<0.05). *(C)* Testing the association between NCS synchrony and the presence of alcohol and food cues. Dynamic inter-subject correlation (ISC) of NCS activation during video-watching was computed for each subject pair using the time series of the NCS (**Figure 2D**). ISC was computed within sliding windows (window size=10 TR, step size=1 TR) to capture fluctuations in NCS synchrony over time. After standardizing each time-series, the dynamic ISC of NCS was correlated with the time-series of alcohol and food cue presence. Data before standardization are shown. *(D)* Results of the correlation analyses. In the line graph, the X-axis represents time, and the Y-axis shows the standardized values of the averaged pairwise dynamic ISC (black), alcohol cue presence (red), and food cue presence (blue). The histograms show null distribution of correlation coefficients obtained from an exhaustive circular shift permutation test (N=700 unique shifts), with the observed correlation coefficient indicated by the black line.

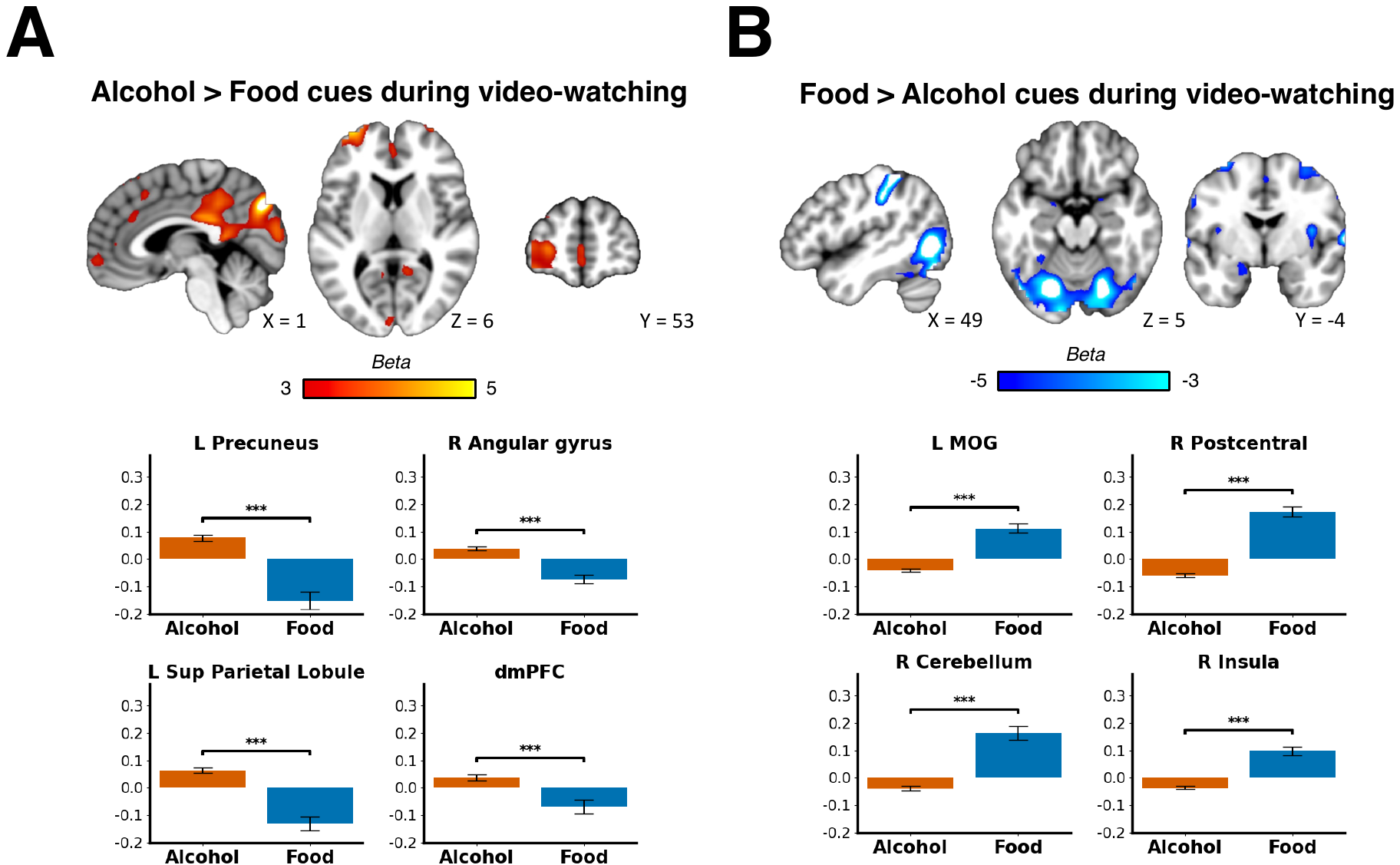

**Supplementary Figure S9.** Whole-brain voxel-wise second-level results comparing activation in response to alcohol versus food cues. *(A)* Voxels showing greater activation to alcohol cues compared to food cues. *(B)* Voxels showing greater activation to food cues compared to alcohol cues. The bar graphs display the mean beta coefficients for alcohol and food contrasts, with error bars representing the standard error of the mean. General linear model (GLM) results were thresholded at *P*<0.05 (FDR corrected) with an extent threshold of *k* ≥ 50 voxels. *Acronyms*: MOG, Medial Occipital Gyrus; dmPFC; Dorsomedial Prefrontal Cortex.

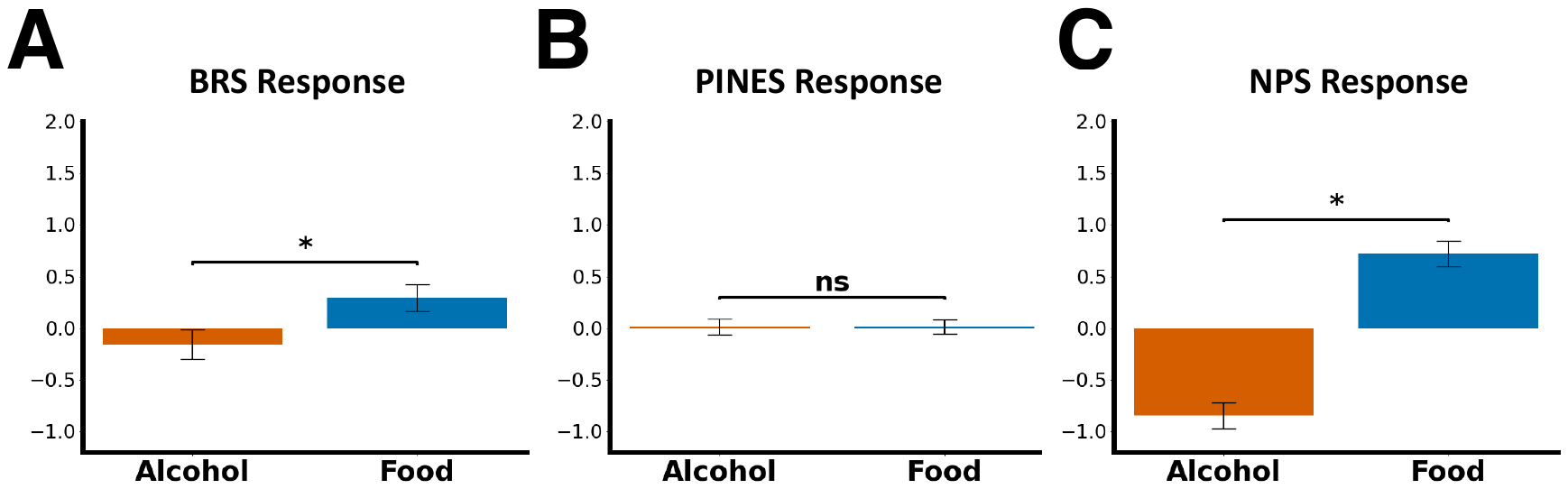

**Supplementary Figure S10.** Comparison of other neuromarkers' activation in response to alcohol and food cues. Comparison of *(A)* Brain Reward Signature (BRS) response, *(B)* Picture-Induced Negative Emotion Signature (PINES) response, and *(C)* Neurologic Pain Signature (NPS) response to alcohol and food cues. Based on the first-level general linear model (GLM) fitted to compare brain responses during the presentation of alcohol and food cues, a single response score for each neuromarker (replacing the neurobiological craving signature NCS) was extracted per participant for each contrast (**Figure S8A**). The X-axis represents the first-level contrast of “alcohol” and “food,” and the Y-axis shows the mean response of each neuromarker, with error bars indicating standard error. A two-sample t-test was performed to compare the responses of each neuromarker to alcohol and food cues. The asterisk indicates a significant difference (*P*<0.05).

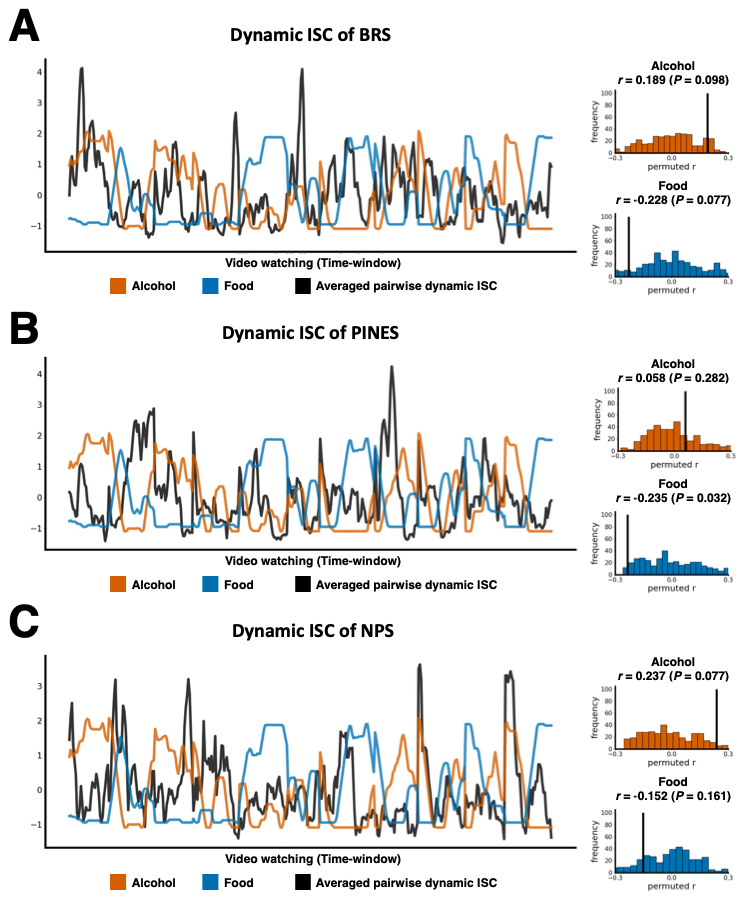

**Supplementary Figure S11.** Correlation between dynamic inter-subject correlation (ISC) of each neuromarker and presence of alcohol and food cues**.** For each neuromarker, we computed the averaged dynamic inter-subject correlation (ISC) across all participant pairs using a sliding window. The resulting time series was z-scored and then correlated with the z-scored time series of cue presence (alcohol and food, respectively). *(A)* Brain Reward Signature (BRS), *(B)* Picture-Induced Negative Emotion Signature (PINES), and *(C)* Neurologic Pain Signature (NPS) each show the dynamic ISC (black), alcohol cue presence (orange), and food cue presence (blue) over time. The histograms show null distribution of correlation coefficients obtained from an exhaustive circular shift permutation test (N=700 unique shifts), with the observed correlation coefficient indicated by the black line.

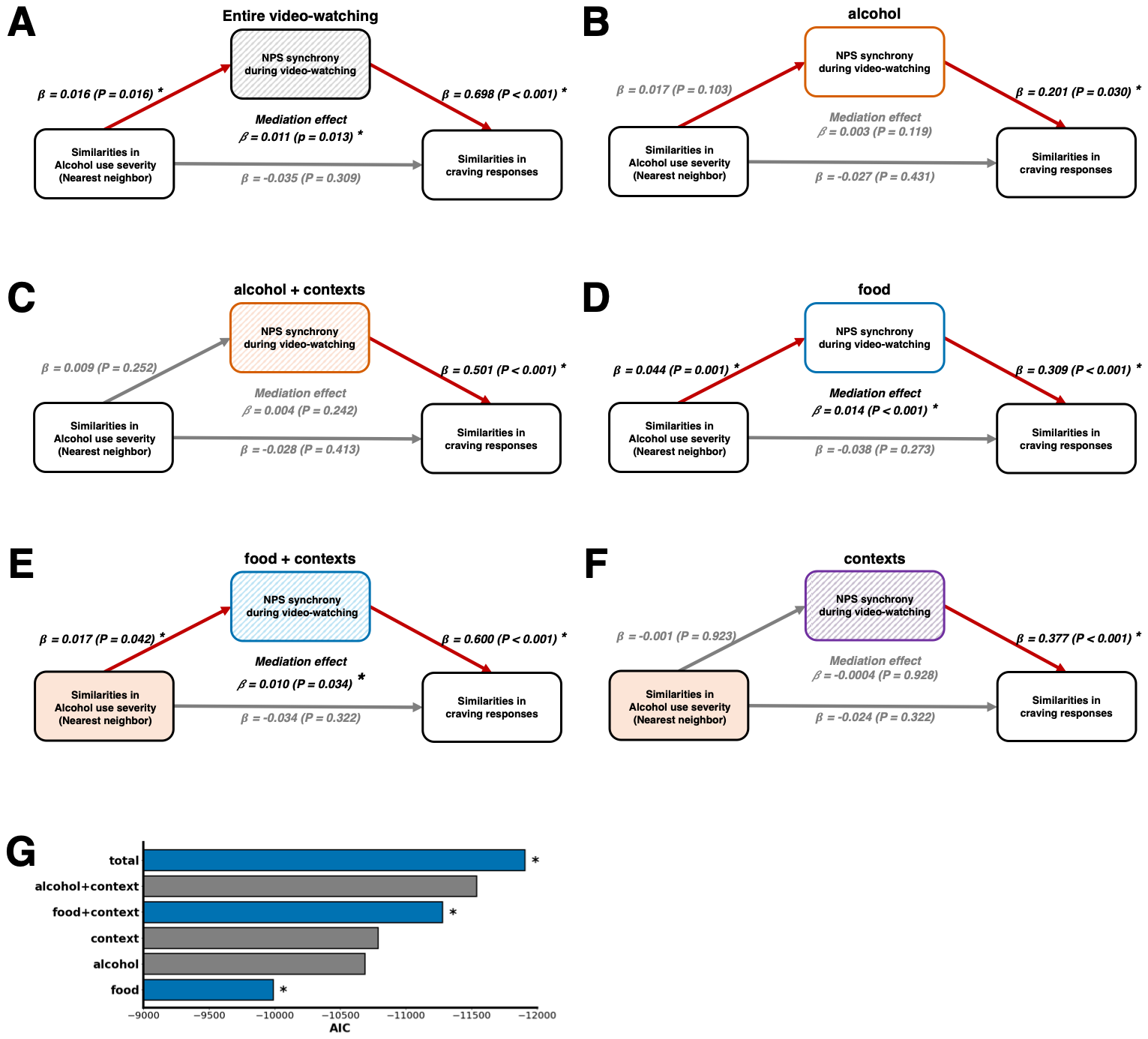

**Supplementary Figure S12.** Mediation models investigating neural synchrony of the Neurologic Pain Signature (NPS) as a mediator. Mediation analyses examined whether the relationship between similar drinking reasons and similar craving responses was mediated by the NPS synchrony in response to *(A)* entire video-watching, *(B)* alcohol cues, *(C)* alcohol cues with contexts, *(D)* food cues, *(E)* food cues with contexts, and (F) contexts alone during video-watching. The coefficient and significance of the mediation effect were assessed by bootstrapping of 10,000 samples, with significant beta coefficients (*P*<0.05) marked by asterisks. *(G)* Model comparisons show Akaike Information Criterion (AIC) values, where lower AIC indicates better fit. Colored bars highlight models with significant mediation effects (*P*<0.05).

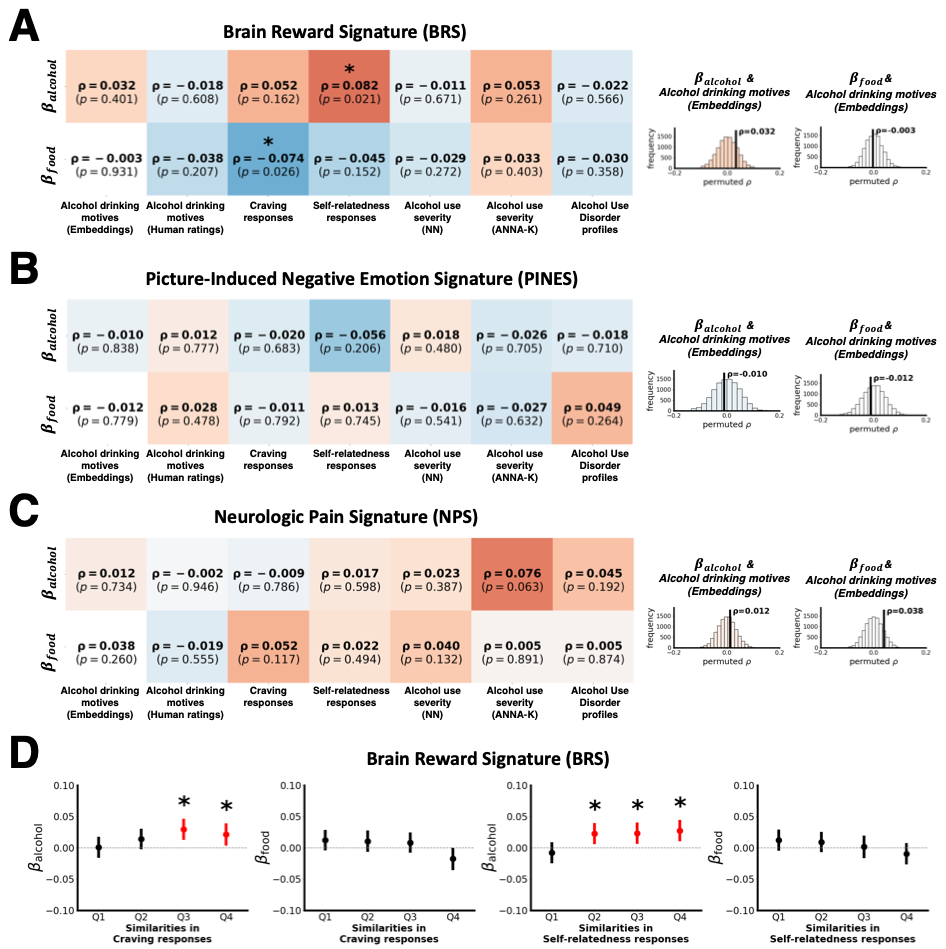

**Supplementary Figure S13.** Regression analysis of dynamic inter-subject correlation (ISC) of other neuromarkers in response to alcohol and food cues. Results of the correlation analyses after running regression analysis (**Figure 3A-B**) using *(A)* Brain Reward Signature (BRS), *(B)* Picture-Induced Negative Emotion Signature (PINES), and *(C)* Neurologic Pain Signature (NPS). Significance was determined via permutation testing with 10,000 permutations. Histograms show the null distribution of permuted coefficients, with the actual correlation coefficient indicated by black vertical lines. Asterisks denote significant beta coefficients (*P*<0.05). *(D)* Analysis of absolute beta coefficients related to similarities in behavioral measures with significant correlation (i.e., craving and self-relatedness responses), focusing on significant relationships involving the BRS. The X-axis represents similarity values divided into four quantiles (Q1: least similar, Q4: most similar), while the Y-axis displays mean beta coefficients for each quantile. Error bars represent 95% confidence intervals. Asterisks indicate statistically significant mean beta coefficients based on one-sample t-tests.

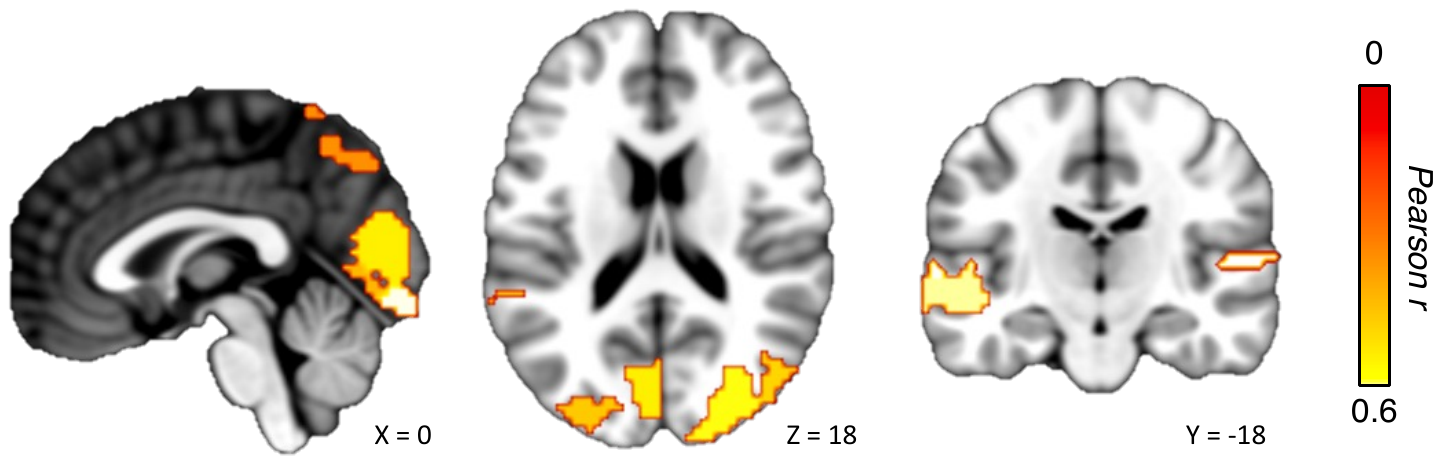

**Supplementary Figure S14.** Inter-subject correlation (ISC) during video-watching. Watching alcohol-drinking videos evokes widespread ISC across the entire sample (N=53). Colored regions of interest (ROIs; Shen et al. (4)) indicate significant synchronized activation across participants. To determine the significance of ISC for each ROI, a non-parametric circular shift randomization with 10,000 permutations was employed, in which shift amounts were randomly sampled with replacement at each permutation to generate a null distribution. ROIs where all participants showed significant leave-one-out ISC (*P<*0.05) were considered significant.

**Supplementary Table S1.** Inter-subject representational similarity analysis (IS-RSA) using neural similarity matrix of the neurobiological craving signature (NCS) during video-watching

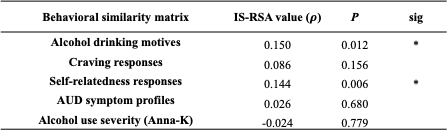

Note. Significance was assessed using permutation testing with 10,000 permutations. Asterisks indicate significant Spearman correlation between the behavioral and the neural similarity matrix (**Figure 2E**). P-values are uncorrected. Acronyms: AUD, Alcohol Use Disorder; Anna-K, Anna-Karenina.

**Supplementary Table S2.** Multiple linear regression explaining the NCS synchrony during video-watching: controlling for other behavioral similarities

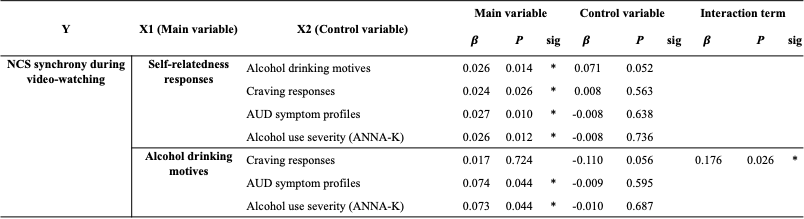

Note. Results of multiple linear regression analyses using pairwise similarity values, with each row displaying the result of a separate model. The dependent variable in all models is the similarity in the response of neurobiological craving signature (NCS) during video-watching. X1 represents the main independent variable of interest, and X2 denotes the control variable. Each row displays the model of $Y \sim X1 + X2$. An interaction term ($X1\cdot X2$) was included in all models but removed if non-significant. For models where the interaction term was significant, only the results including the interaction are shown. The significance of each beta coefficient was determined based on permutation testing with 10,000 permutations. P-values are uncorrected. Asterisks indicate significant beta coefficients. Acronyms: AUD, Alcohol Use Disorder; Anna-K, Anna-Karenina.

**Supplementary Table S3.** Multiple linear regression explaining the NCS synchrony during video-watching: controlling for other speech embedding vectors

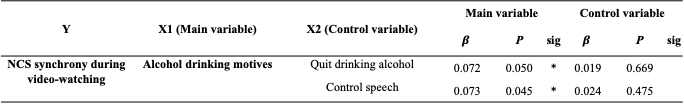

Note. Results of multiple linear regression analyses using pairwise similarity values. Each row represents a separate model in which the dependent variable is the similarity in the Neurobiological Craving Signature (NCS) during video-watching. X1 denotes the main independent variable of interest: similarity in alcohol drinking motives (speech embedding vectors). X2 represents the control variable: similarity in speech about quitting drinking alcohol (i.e., in response to the prompt “Please share your thoughts about quitting, refraining, or reducing drinking alcohol.”) and control speech (i.e., in response to the prompt “Please share what happened before you came to this laboratory today.”), both processed into speech embedding vectors. Each row displays the model of $Y \sim X1 + X2$. An interaction term ($X1\cdot X2$) was included in all models but removed if non-significant. The significance of each beta coefficient was determined based on permutation testing with 10,000 permutations. Asterisks indicate significant beta coefficient (P<0.05).

**Supplementary Table S4.** Inter-subject representational similarity analysis (IS-RSA) using neural similarity matrix of the Brain Reward Signature (BRS) during video-watching

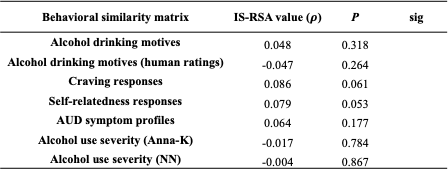

*Note.* Significance was assessed using permutation testing with 10,000 permutations. P-values are uncorrected. Acronyms: AUD, Alcohol Use Disorder; NN, Nearest-neighbor; Anna-K, Anna-Karenina.

**Supplementary Table S5.** Inter-subject representational similarity analysis (IS-RSA) using neural similarity matrix of the Picture-Induced Negative Emotion Signature (PINES) during video-watching

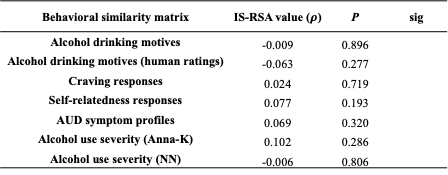

*Note.* Significance was assessed using permutation testing with 10,000 permutations. P-values are uncorrected. Acronyms: AUD, Alcohol Use Disorder; NN, Nearest-neighbor; Anna-K, Anna-Karenina.

**Supplementary Table S6.** Inter-subject representational similarity analysis (IS-RSA) using neural similarity matrix of the Neurologic Pain Signature (NPS) during video-watching

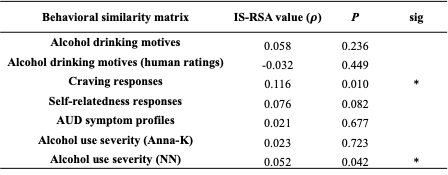

*Note.* Significance was assessed using permutation testing with 10,000 permutations. P-values are uncorrected. Acronyms: AUD, Alcohol Use Disorder; NN, Nearest-neighbor; Anna-K, Anna-Karenina.

**Supplementary Table S7.** Multiple linear regression explaining the NPS synchrony during video-watching: controlling for other behavioral similarities

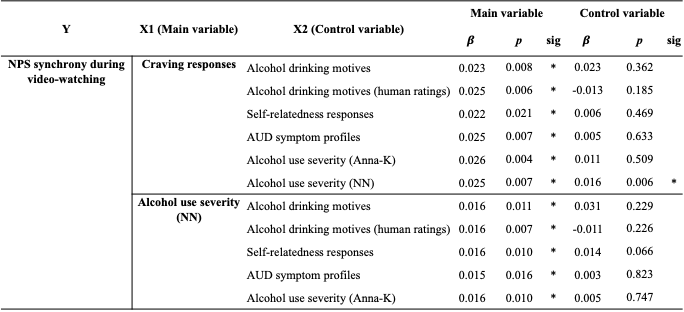

*Note.* Results of multiple linear regression analyses using pairwise similarity values, with each row displaying the result of a separate model. The dependent variable in all models is the similarity in the response of the Neurologic Pain Signature (NPS) during video-watching. X1 represents the main independent variable of interest, and X2 denotes the control variable. Each row displays the model of $Y \sim X1 + X2$. An interaction term ($X1\cdot X2$) was included in all models but removed if non-significant. The significance of each beta coefficient was determined based on permutation testing with 10,000 permutations. P-values are uncorrected. Asterisks indicate significant beta coefficients. Acronyms: AUD, Alcohol Use Disorder; NN, Nearest-neighbor; Anna-K, Anna-Karenina.

1. Park, Eunjeong L., & Cho, Sungzoon. (2014). KoNLPy: Korean natural language processing in Python. Proceedings of the 26th Annual Conference on Human & Cognitive Language Technology, Chuncheon, Korea. KoNLPy Documentation
2. Park, D., Jang, Y., & Kim, H. (2021). Korean-English Machine Translation with Multiple Tokenization Strategy. Proceedings of KCC 2021, Korean Institute of Information Scientists and Engineers (KIISE).
3. Park, D., Jang, Y., & Kim, H. (2021). Korean-English Machine Translation with Multiple Tokenization Strategy. Proceedings of KCC 2021, Korean Institute of Information Scientists and Engineers (KIISE).
4. Reimers, N., & Gurevych, I. (2019). Sentence-BERT: Sentence embeddings using Siamese BERT-networks. Proceedings of the 2019 Conference on Empirical Methods in Natural Language Processing (EMNLP). Retrieved from https://www.sbert.net
5. Monologg. (2020). KoBERT: Korean BERT pre-trained language model. Hugging Face. Retrieved from https://huggingface.co/monologg/kobert
6. Kim, K. (2020). Pretrained Language Models For Korean. GitHub. Retrieved from https://github.com/kiyoungkim1/LMkor
